# Structural and molecular basis of cross-seeding barriers in amyloids

**DOI:** 10.1101/2020.07.06.188508

**Authors:** A. Daskalov, D. Martinez, V. Coustou, N. El Mammeri, M. Berbon, L.B. Andreas, B. Bardiaux, J. Stanek, A. Noubhani, B. Kauffmann, J.S. Wall, G. Pintacuda, S.J. Saupe, B. Habenstein, A. Loquet

## Abstract

Neurodegenerative disorders are frequently associated with β-sheet-rich amyloid deposits. Amyloid-forming proteins can aggregate under different structural conformations known as strains, which can exhibit a prion-like behaviour and distinct patho-phenotypes. Precise molecular determinants defining strain specificity and cross-strain interactions (cross-seeding) are currently unknown. The HET-s prion protein from the fungus *Podospora anserina* represents a model system to study the fundamental properties of prion amyloids. Here, we report the amyloid prion structure of HELLF, a distant homolog of the model prion HET-s. We find that these two amyloids, sharing only 17% sequence identity, have nearly identical β-solenoid folds but lack cross-seeding ability *in vivo*, indicating that prion specificity can differ in extremely similar amyloid folds. We engineer the HELLF sequence to explore the limits of the sequence-to-fold conservation and to pinpoint determinants of cross-seeding and prion specificity. We find that amyloid fold conservation occurs even at an exceedingly low level of identity to HET-s (5%). Next, we derive a HELLF-based sequence, termed HEC, able to breach the cross-seeding barrier *in vivo* between HELLF and HET-s, unveiling determinants controlling cross-seeding at residue level. These findings show that virtually identical amyloid backbone structures might not be sufficient for cross-seeding and that critical side-chain positions could determine the seeding specificity of an amyloid fold. Our work redefines the conceptual boundaries of prion strain and shed new light on key molecular features concerning an important class of pathogenic agents.

## Introduction

Amyloid-forming proteins undergo a phase transition to form insoluble, polymeric assemblies, which can self-propagate *in vivo* as prions (1–3). Amyloid aggregates associated with neurodegenerative diseases (i.e. Alzheimer’s, Parkinson’s etc.) have a prion-like behaviour (4, 5) and can propagate under different structural conformations also known as prion ‘strains’, which may be associated with distinct phenotypes of the pathology (6–9). The cross-talk between an infectious amyloid conformation (prion strain) and a naive homologous or heterologous amyloidogenic sequence, referred to as ‘cross-seeding’ (10, 11), is a critical event in prion biology, representing the key aspect of infectivity. Amyloid cross-seeding could play a role in the aetiology (12) and pathogenesis of various proteinopathies (13, 14). However, our understanding of the precise molecular and structural determinants allowing or limiting cross-seeding remains poor.

The fungal HET-s protein constitutes a highly favourable system to study the fundamental properties of prion amyloids, as a high-resolution structure of the propagative prion state is available (15, 16). The prion-forming domain (PFD) of HET-s contains two 21 amino acid pseudo-repeats (R1 and R2), which are alternately stacked, each repeat forming four β-strands, adopting a left-handed β-solenoid fold (15, 16). Noteworthy, several reports have stressed structural similarities between the HET-s amyloid fold and pathological amyloids formed by the human prion protein PrP(17) and tau (18).

Despite the structural similarities with some pathological amyloids, HET-s represents a functional amyloid, which is integral to an immunity-related signal transduction pathway in fungi (19). The β-solenoid fold ensures signal transduction from an activated NOD-like receptor (NLR) to a downstream execution protein (HET-S, a pore-forming variant of the HET-s prion protein). The fold represents a cell death trigger and functions on the basis of the prion principle (19, 20). The structural templating of the HET-S PFD into the amyloid fold triggers the cytotoxicity of the α-helical HeLo domain, which induces cell death targeting the plasma membrane (21–23). Signal-transducing amyloids are widespread in fungi (24) and at least five subfamilies of HET-s-related amyloid motifs (HRAMs) have been identified (25). The HRAMs exhibit the two pseudo-repeats organization of HET-s and a pattern of hydrophobic and polar amino acids, while each subfamily is being defined by the conservation of a set of HRAM-specific residues (25). It has been speculated that the HRAMs may share a common β-solenoid fold and that natural diversification is driven to preserve specificity of the signalling pathways (between NLRs and cognate effectors) by limiting amyloid cross-seeding between distinct HRAMs (19, 25). Here, we use the HRAMs as an experimental framework to investigate 1) the sequence-to-fold relation in prion amyloids and 2) the molecular determinants defining prion specificity, which allow or prevent cross-seeding.

## Results

### Molecular characterization of HELLF

We chose to work with a newly identified HELLF protein, encoded in the genome of *Podospora anserina* (*SI Appendix*, Fig. S1), the natural host of the [Het-s] prion. HELLF consists of an N-terminal HELL (HeLo-like) domain and a putative C-terminal PFD. The HELLF PFD carries strongly divergent HRAM pseudo-repeats (R1 and R2) showing only 17% of sequence identity with the PFD repeats of HET-S/s (Fig. 1A; *SI Appendix*, Fig. S1). HELLF pseudo-repeats belong to the HRAM5 family (Fig. 1B). Strains expressing HELLF(209-277) showed phenotypic bistability exhibiting either a [Φ*] phenotype, where the protein remains soluble or a [Φ] phenotype with formation of dot-like aggregates (Fig. 1C; *SI Appendix*, Fig. S1). The [Φ] strains triggered cell death upon anastomosis (cellular fusion) with strains expressing full-length HELLF, while [Φ*] strains formed viable heterokaryons (Fig. 1D). During the cell death reaction, as described for HET-S (22), HELLF relocates to the cell membrane (*SI Appendix*, Fig. S1). The [Φ*] strains (aggregates-free) switched spontaneously (~24 hours) to [Φ] strains (with aggregates), which in turn could be cured (or reversed) back to [Φ*] state through a sexual cross (*SI Appendix*, Table S1, Table S2). These results establish HELLF as a distant HET-S homolog and demonstrate that the HRAM5 pseudo-repeats bearing PFD of HELLF behaves as a prion in *P. anserina*.

**Fig. 1.**
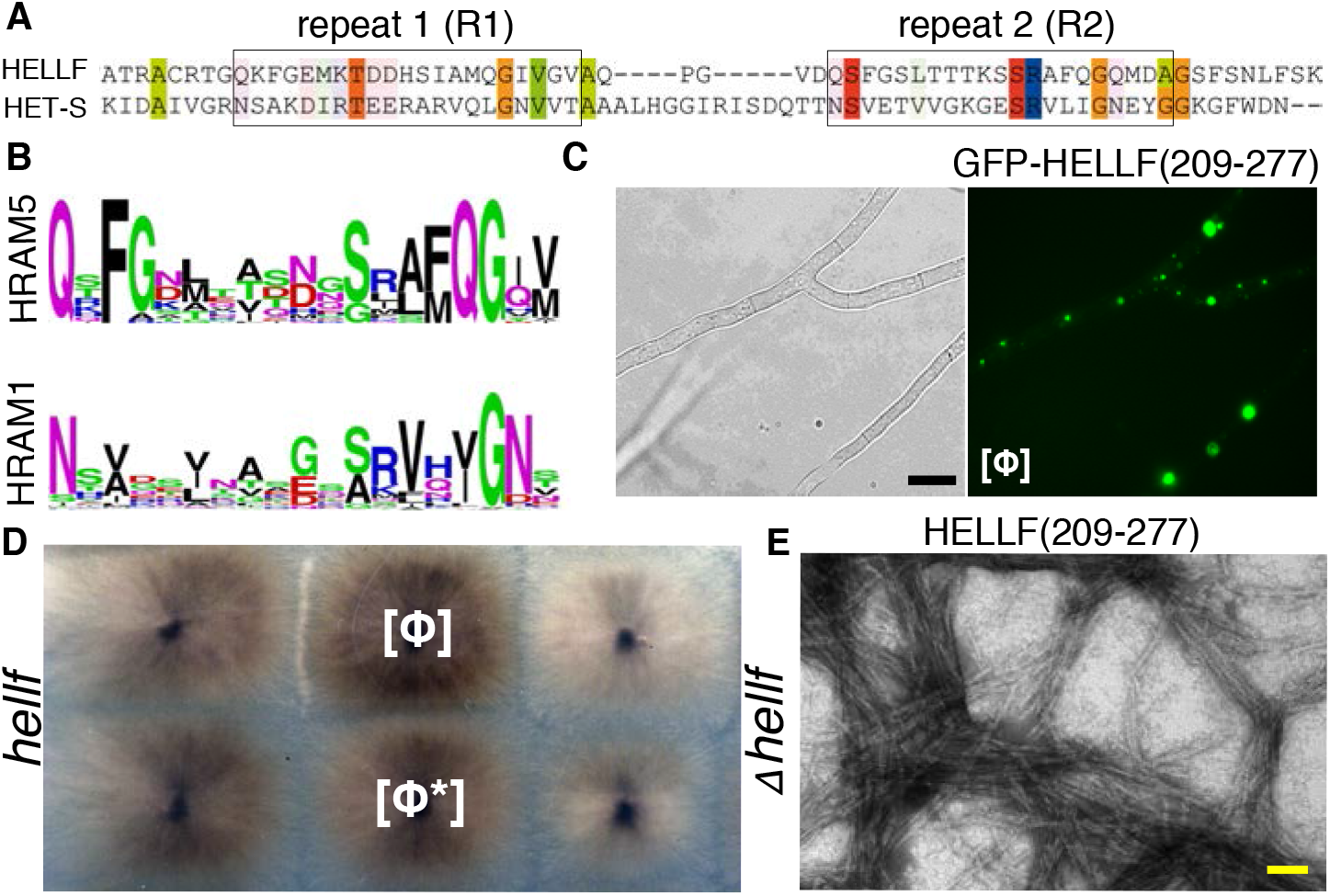
HELLF is a distant HET-s homolog encoded in the genome of *Podospora anserina*. **A.** Sequence homology between HELLF and HET-s PFDs. The two pseudo-repeats are boxed. Conserved residues are shown in Taylor colour scheme and intensity of colour reflects degree of conservation. **B.** MEME cartoons showing the conservation of HRAM-specific residues. **C.** Dot-like aggregates of the HELLF PFD (GFP-HELLF(209-277)) formed in *P. anserina*. Scale bar is 5 μm. **D.** HELLF controls a programmed cell death reaction as determined by the ‘barrage’ phenotype (white arrowhead) between strains expressing full-length HELLF and strains expressing the HELLF PFD in the [φ] prion state. **E.** Electron micrograph of fibrillar assemblies of the HELLF PFD, scale: 100nm.

### Solid-state NMR (ssNMR) structure of HELLF prion domain

We engaged in the structural characterization of HELLF(209-277). The protein self-assembled into unbranched fibrils *in vitro* (Fig. 1E), exhibiting a typical cross-β signature by X-ray diffraction (*SI Appendix*, Fig. S2). We took advantage of recent developments for fast magic-angle spinning (MAS) NMR probes (26–28) to establish, and implement for the first time on a fibril sample, a three-dimensional (3D) structure determination approach, based entirely on ^1^H-^1^H proximities. The approach, reminiscent of nuclear Overhauser effect (nOe)-based solution NMR methods to solve soluble globular protein structures, allows a tremendous gain in sensitivity and enables the use of simplified labelling schemes and minimal sample quantities.

Approximately 300μg of fully protonated HELLF(209-277) were packed into a 0.7 mm ssNMR rotor (Fig. 2A). Ultra-fast MAS performed at rates of ~110 kHz allowed the acquisition of high-resolution !H-detected multidimensional spectra even in fully protonated samples (Fig. 2B; *SI Appendix*, Fig. S3) with ^1^H line widths of ~150-200Hz enabling assignment of backbone and side chain protons (29). A unique set of resonances revealed the presence of a single conformational polymorph in the fibrillar assembly, and conformation-dependent chemical shifts (30) revealed a β-rich rigid core extending from residue Q221 to S272, including 8 β-strands (Fig. S4). A flexible linker segment (G240-D247), comprising a β-breaker GxxxPG motif, subdivides the rigid core in 2 regions with 4 β-strands each (Fig. S4). We employed a combination of 3D H(H)CH and H(H)NH experiments (27) on a fully protonated, uniformly ^13^C,^15^N-labeled sample to derive 178 internuclear ^1^H-^1^H distances (Fig. 2C). 3D amyloid architectures render the distinction between intra- and intermolecular contacts in ssNMR experiments difficult and usually require complex labelling schemes (16, 31). We designed a new and simple labelling strategy based on an equimolar mixture of fully protonated proteins at natural abundance, randomly co-aggregated with deuterated, extensively amide-reprotonated and uniformly ^15^N-labeled proteins (scheme denoted as (1/1) [(U-^1^H,^14^N)/(U-^1^H^N^,^2^H,^15^N)]) (Fig. 2E). Using this scheme, we observed 33 intermolecular ^1^H-^1^H inter-strand aliphatic to amide distances relying on a single 3D H(H)NH spectrum (Fig. 2D) with an asymmetric polarization transfer (Fig. 2F). We identified 211 distance restraints in total, of which 176 are long-range restraints (|i-j| > 4) (Fig. 2G) to derive a 3D structure of the HELLF fibrillar assembly at atomic resolution with 20-conformer bundle r.m.s.d of 0.73Å for backbone atoms and 1.18Å for heavy atoms (Fig. 3; *SI Appendix*, Table S3). Our 3D structure determination approach based on ^1^H-^1^H proximities compares favourably to benchmark studies of the HET-s prion domain amyloid structure by ssNMR (16); we detected approximately 4.5 structurally meaningful restraints per residue in the rigid amyloid core, a number approaching those used in high-resolution structure determination protocols in solution NMR.

**Fig. 2.**
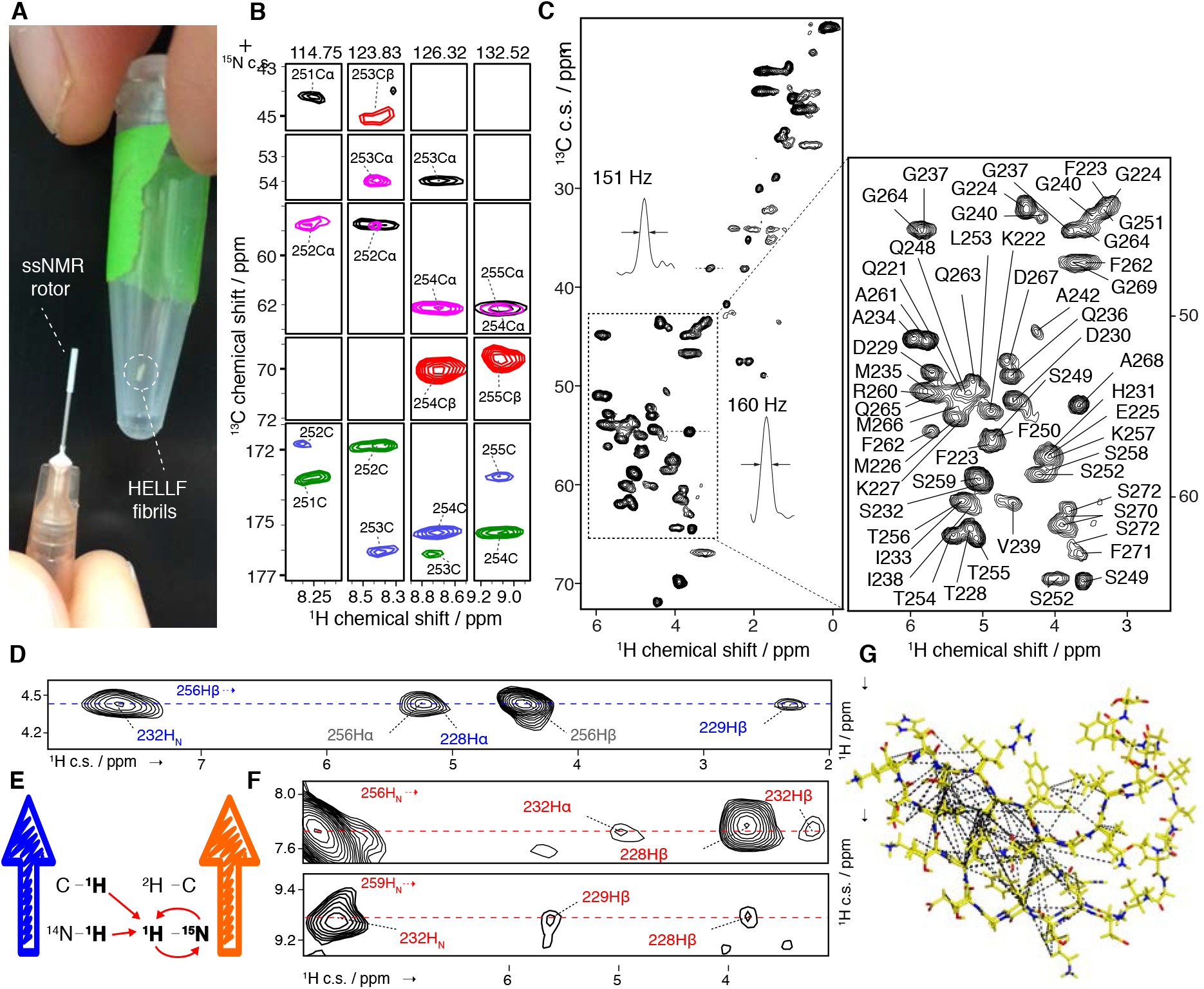
Solid-state NMR (ssNMR) characterization of HELLF(209-277) fibrils. **A.** ssNMR sample preparation of HELLF(209-277), requiring minimal sample quantity (<300ug) using a 0.7 mm ssNMR rotor. **B.** Extracts of ssNMR spectra for ^1^H, ^13^C and ^15^N sequential assignments. A combination of (HCA)CB(CA)NH (red), (HCO)CA(CO)NH (black), (H)CANH (purple), (H)CONH (green) and (H)CO(CA)NH (blue) was used to assign HELLF(209-277) fibrils. **C.** ^1^H-^13^C ssNMR spectra of fully protonated HELLF(209-277) amyloid fibrils. **D-F.** Collection of ssNMR distance restraints. **D.** Intramolecular distances based on a H…(H)CH experiment on fully protonated HELLF sample. **E.** ssNMR approach to detect intermolecular interactions, based on a (1/1) [(U-^1^H/^14^N)/(U-^1^H_N_/^2^H/^15^N)]-labelled HELLF sample. **F.** Intermolecular distances based on a H.(H)NH experiment on (1/1) [(U-^1^H/^14^N)/(U-^1^H_N_/^2^H/^15^N)]-labelled HELLF(209-277). **G.** ssNMR restraints shown on the HELLF(209-277) rigid core.

**Fig. 3.**
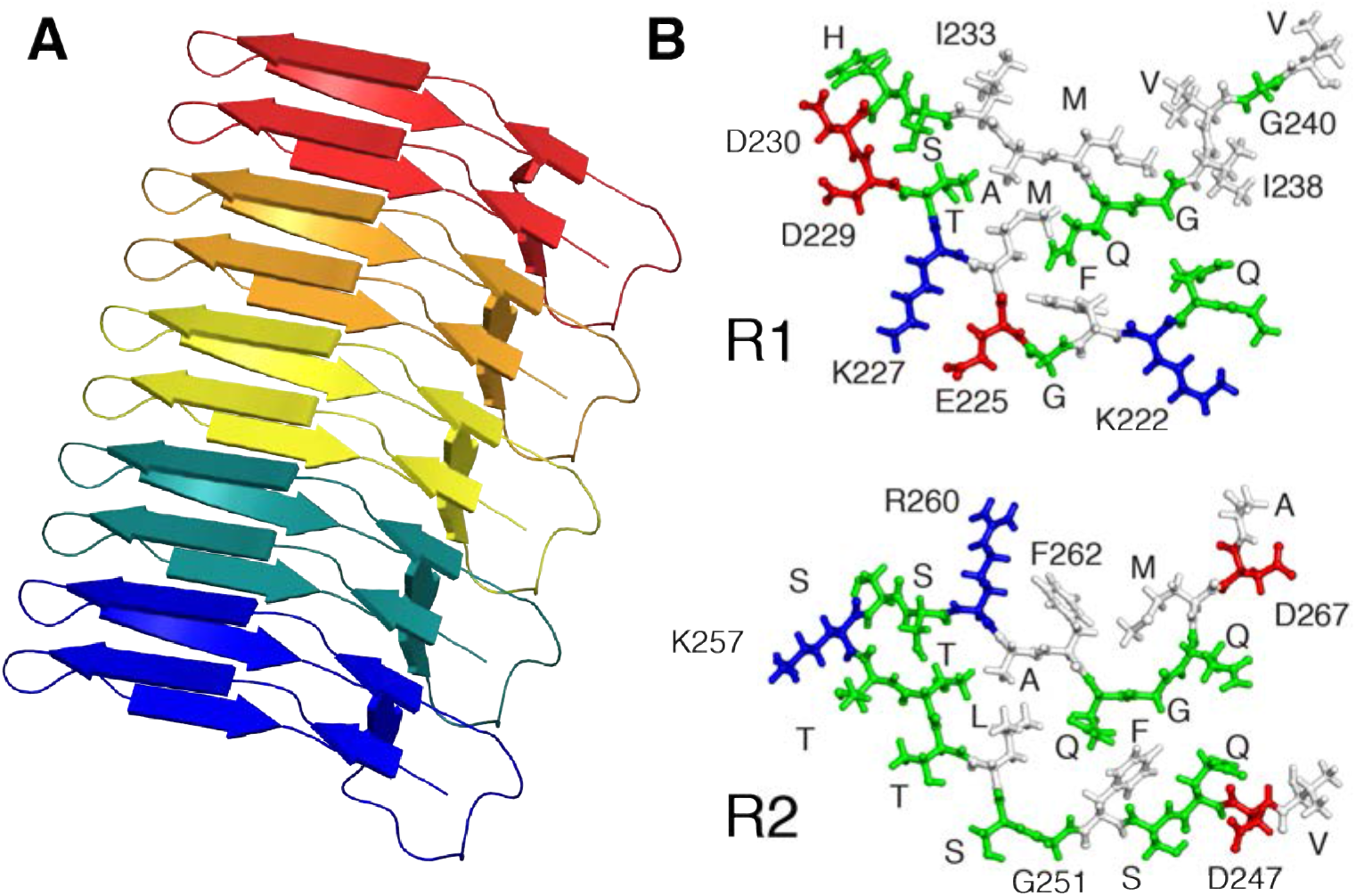
Solid-state NMR structure of HELLF(209-277) amyloid assembly. **A.** Side view of a ribbon representation of HELLF(209-277) structure, representing five monomers stacked in the fibrillar arrangement. Individual molecules are represented in different colours. **B.** Top view of the solenoid-forming pseudo-repeats, R_1_ (top) and R_2_ (bottom), of HELLF(209-277). Hydrophobic residues are shown in white, acidic residues in red, basic residues in blue and others green.

### HELLF and HET-s prions share an identical backbone β-solenoid core

The HELLF fibrillar architecture shows a β-solenoid fold, made by the intermolecular packing of HELLF monomers along the fibril axis (Fig. 3). The intramolecular HELLF fold is composed of 8 β-strand structural elements (β1 to β8) separated by a short unstructured region (Gly240-Asp247) (Fig. 3). Intramolecular ssNMR restraints reveal a regular and rigid core, constituted by alternate stacking of R1 (β1 to β4) and R2 (β5 to β8) pseudo-repeat regions (Fig. 4B, Fig. 3). Each pseudo-repeat adopts a triangular shape, stabilized by hydrophobic side-chains interlaced inside the amyloid core and protected from the solvent. Several β-breaker glycine residues allow for bending two consecutive β-arcs into the triangular arrangement of the amyloid core (Fig. 4B). R1 and R2 repeats are stacked through a hydrogen bond-rich pairing between β-sheets (β_i_ with β_i+4_, T255/S232 and T228/S259). To corroborate the HELLF intramolecular R1/R2 stacking, we performed scanning transmission electron microscopy (STEM) mass-per-length (MPL) measurements and determined a MPL of O.99±O.1O kDa/Å corresponding to 1.1±0.1 molecules per 0.94 nm (*SI Appendix*, Fig. S5). Considering a β-strand repetition of 0.47 nm in the cross-β architecture as measured by X-ray diffraction (*SI Appendix*, Fig. S2), it leads to a fibril layer (i.e. per 0.47 nm) composed of half of a HELLF molecule, in agreement with the fold determined by ssNMR (Fig. 2G, Fig. 3). The inter-subunit packing consists of parallel, pseudo in-register stacking and the overall intermolecular arrangement is consistent with cross-β stacked solenoid architecture. In spite of the low sequence identity between HET-s and HELLF PFDs, both prion domains adopt virtually identical backbone conformations (Fig. 4, A and C).

**Fig. 4.**
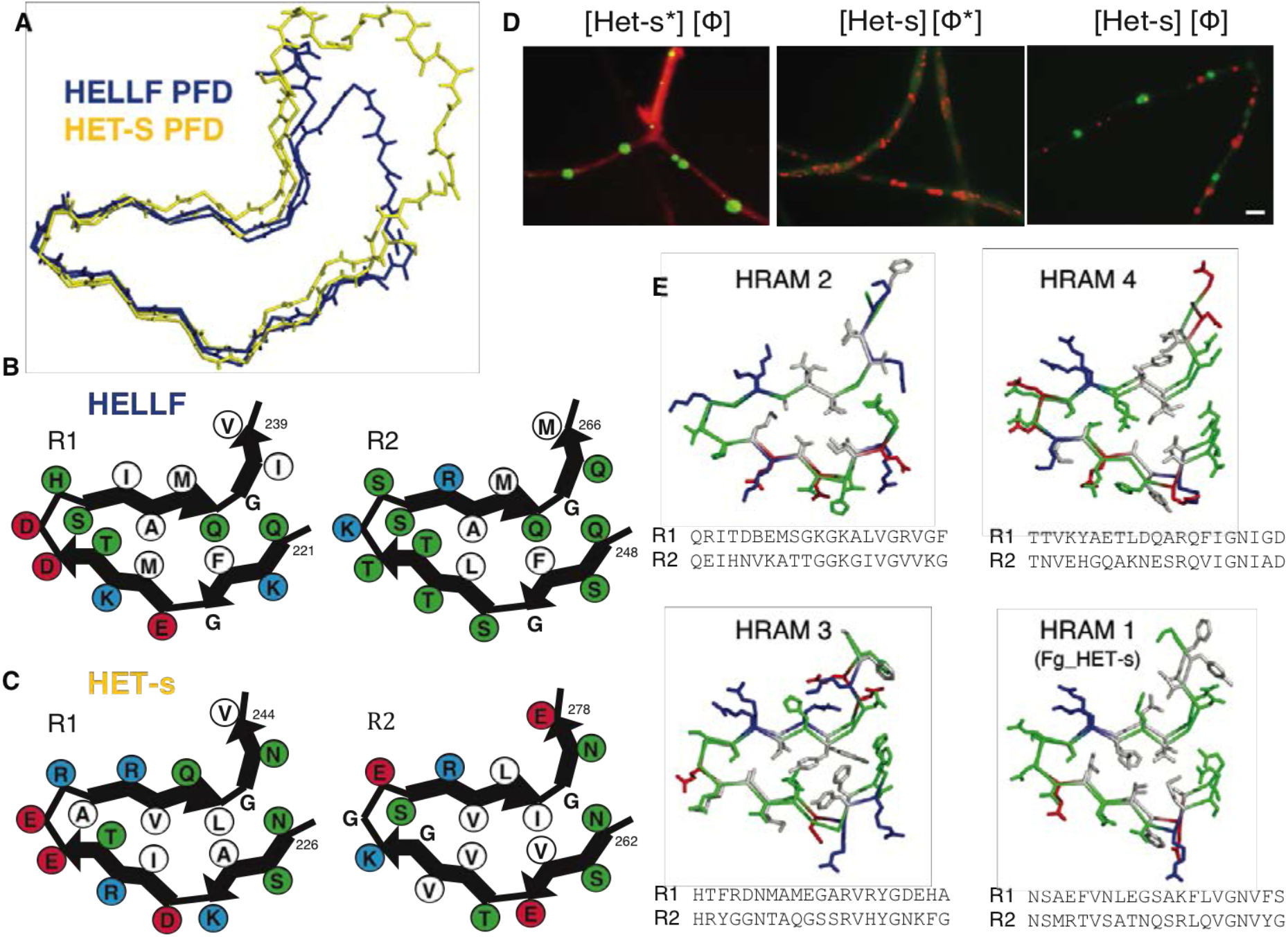
HELLF(209-277) amyloid fold presents strong similarity with HET-s(218-289), while the two prion domains lack *in vivo* cross-seeding. **A.** Backbone structural alignment of HELLF PFD (in blue) and HET-s PFD (in yellow) ssNMR structures. **B-C.** Shown are four cartoons representing the hydrophobic triangular core of amyloid fibrils formed by the successively stacked pseudo-repeats – R1 (left panels) and R2 (right panels) - of HELLF (**B**) and of HET-s (**C**). Amino acid residues decorating the amyloid backbones are drawn as beads of different colours. Hydrophobic residues are shown in white, acidic residues in red, basic residues in blue and others green. **D.** Fluorescent microscopy images of strains co-expressing HET-s-RFP ([Het-s*] or [Het-s] state) and the cytotoxic-dead HELLF(L52K)-GFP mutant ([Φ*] or [Φ] state) exhibiting distinct epigenetic combinations. Full panels are given in supplementary Figure S8. Scale bar: 2 μm. **E.** 3D models for the two pseudo-repeats of different HRAM families.

We took advantage of the high-resolution structures of HELLF and HET-s to perform molecular modelling on the remaining HRAM families identified in fungal genomes (Fig. 4E). We found that the observed sequence diversity in the HRAM superfamily is indeed compatible with a unique structural solution underlying this amyloid fold (Fig. 4E). In addition, we designed a synthetic protein sequence termed HED (HET-s distant) carrying two identical repeats that share less than 5% identity with HET-s (one residue from 21) (*SI Appendix*; Fig. S6, Table S5). In spite of the extremely low sequence identity to HET-s, HED was able to form a prion *in vivo* and to adopt a HET-s-like β-solenoid structure *in vitro* (*SI Appendix*, Fig. S6).

### HELLF and HET-s carry distinct prion specificities *in vivo*

Considering the structural similarity between HELLF and HET-s prion folds (Fig. 4A) and that these proteins occur in the same species, we analysed the cross-seeding of [Φ] and [Het-s] prion states *in vivo* (*SI Appendix*, Table S2). We found that [Φ] strains do not convert non-prion [Het-s*] strains to the [Het-s] prion state nor do [Het-s] strains induce [Φ] prion formation (*SI Appendix*, Table S2). HELLF PFD fibrils show no [Het-s] infectivity and HET-s PFD fibrils show no [Φ] infectivity in transfection assays (*SI Appendix*, Table S4). HELLF and HET-s PFDs form independent aggregates *in vivo* indicating that the two prions do not co-aggregate (Fig. 4D; *SI Appendix*, Fig. S7). Strains co-expressing HELLF and HET-s can display four alternate epigenetic states ([Het-s*] [Φ*], [Het-s] [Φ] but also [Het-s*] [Φ] and [Het-s] [Φ*]) confirming that [Het-s] and [Φ] are independent prions (*SI Appendix*, Fig. S7, Table S2). Since the backbone structures in the amyloid fold of HET-s and HELLF PFDs are nearly identical (Fig. 4A), it appears that strong structural similarity and interactions of the cross-β backbones are insufficient for amyloid templating and/or co-aggregation.

### HRAM-specific residues control cross-seeding between HELLF and HET-s

The results suggested that side-chain residues, decorating the amyloid backbone, could play a key role in defining prion specificity and cross-seeding. Thus, we decided to test whether by varying HRAM-specific residues between HET-s and HELLF PFDs, the cross-seeding barrier between the two prions could be breached. We replaced five residues on each of the two 21 amino acids HRAM5 pseudo-repeats of HELLF with residues from the corresponding positions found in the HRAM1 pseudo-repeats of HET-s. The engineered sequence was termed HEC (HET-s closer). We targeted for replacement residues that are essential for distinguishing HRAM5 (HELLF) from HRAM1 (HET-s) (25) (Fig. 5A). The five amino acid substitutions for each pseudorepeat of HELLF were introduced in two strongly conserved HRAM5-specific sub-motifs of the protein - the QxFG (position 1-4, where x is any possible amino acid residue) and QG(QI) (position 16-18) sub-motifs (Fig. 5A, Fig. 1B).

**Fig. 5.**
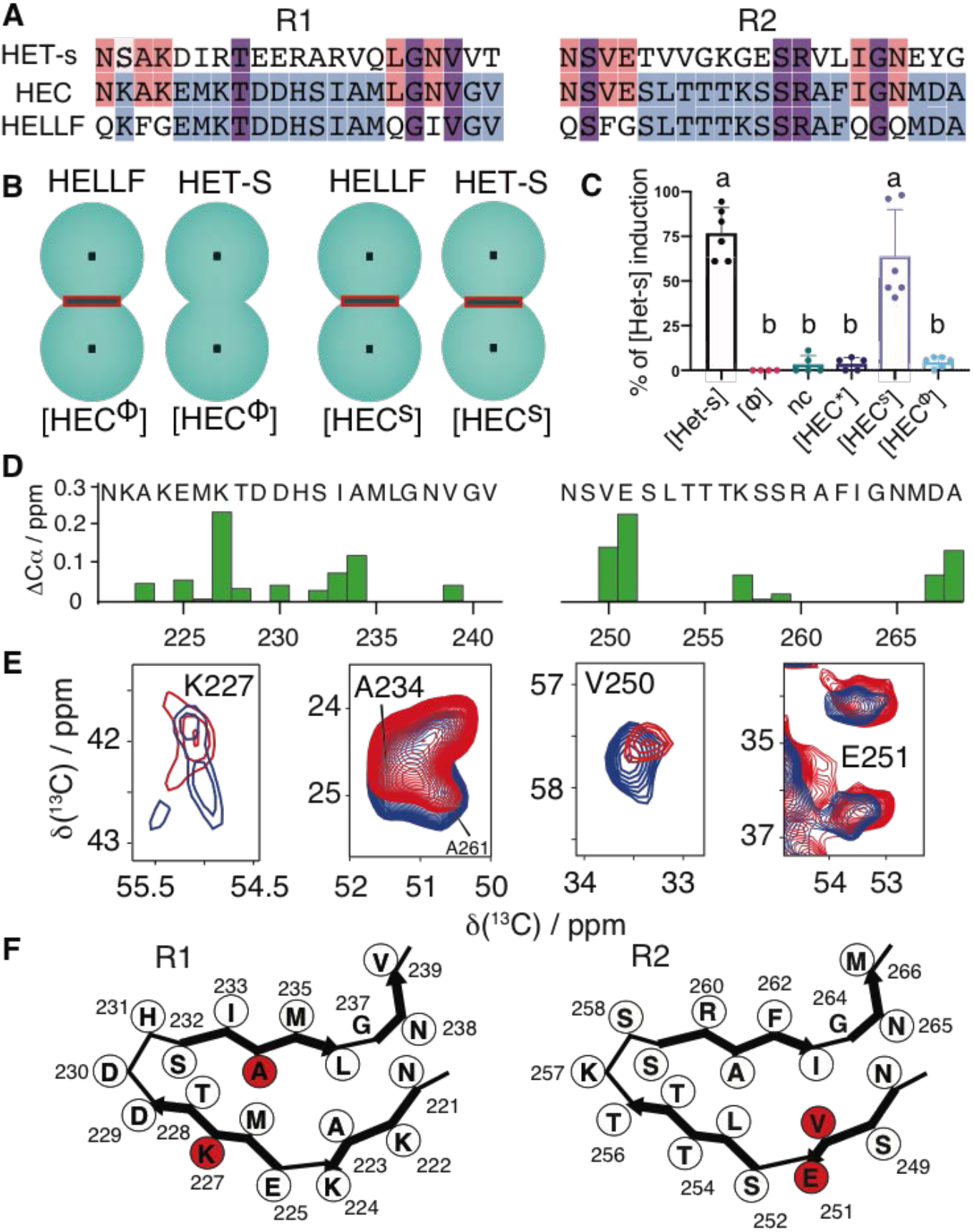
HELLF-derived chimeric protein HEC breaches the prion cross-seeding barrier between HELLF and HET-s. **A.** Sequence alignments of the two pseudo-repeats (R1 and R2), constituting the PFDs of HELLF, HET-s and the engineered protein HEC. HELLF-specific residues are shown in blue colour. HET-s-specific HRAM1-defining residues are shown in red. Residues shared between all three proteins are shown in purple. **B.** Representation of barrage phenotypes between strains expressing full-length HET-S or HELLF in confrontations with HEC-expressing strains of [HEC^S^] (carrying HEC prion strain induced by contact with [Het-s] prion) or [HECΦ] (HEC prion strain induced by [Φ]) phenotypes. The barrage reaction is shown as a line separating two incompatible strains (green circles). **C.** Induction of the [Het-s] prion by HEC-expressing strains *in vivo*. [Het-s] induction is measured in percentage of prion-free [Het-s*] strains converted (by the cited strains) to prion-infected [Het-s] strains. Negative control is indicated with a minus sign. **D.** Cα chemical shift differences between HEC seeded with [Het-s] and [Φ]. **E.** Extracts of ^13^C-^13^C ssNMR spectra of HEC fibrils seeded with [Het-s] (red) or [Φ] (blue). **F.** Cartoon representation of the amyloid backbone of HEC pseudo-repeats. Residues with highest conformational changes between HEC seed by HET-s or HELLF, as measured by difference in chemical shift on Cα, are shown in red.

We expressed HEC in a *Δhet-sΔhellf* strain and tested the ability of the protein to propagate as a prion and to carry [Het-s] and/or [Φ] prion specificity *in vivo*. Strains expressing GFP-HEC showed two distinct phenotypes; [HEC*] strains presented diffuse GFP-HEC fluorescence, did not induce [Het-s] or [Φ] prions, and were unable to trigger cell death by incompatibility with HET-S or HELLF expressing strains (*SI Appendix*, Fig. S8, Table S6, Table S7). We observed that some [HEC*] strains did, spontaneously and at low rate (~10%), transition into a [HEC] state characterized with the appearance of fluorescent dot-like aggregates (*SI Appendix*, Fig. S8, Table S6). None of the spontaneous [HEC] strains were able to induce [Het-s] or produce a barrage with a HET-S strain. Yet, [HEC] strains induced cell death with HELLF-expressing strains, indicating that the engineered HEC sequence shows [Φ] prion specificity and we termed the phenotype [HEC^Φ^] (Fig. 4B; *SI Appendix*, Fig. S8). We were equally able to induce the [HEC^Φ^] phenotype by exposing [HEC*] strains to [Φ] strains (*SI Appendix*, Table S7). Spontaneously formed (or [Φ]-induced) [HEC^Φ^] strains converted [HEC*] strains into [HEC^Φ^] prion state after contact, demonstrating the ability of aggregated HEC to self-propagate (behave as prion) (*SI Appendix*, Fig. S8).

Next, we found that after a contact with a [Het-s] strain, the prion-free [HEC*] strains could equally be converted to a [HEC] state, characterized by the appearance of fluorescent GFP-HEC dot-like aggregates and the ability to induce cell death with HELLF, indicating that the prion state of HEC can be induced both by HELLF and HET-s (*SI Appendix*, Table S7). Importantly, we found that [Het-s]-induced [HEC] strains, unlike the [HEC^Φ^] strains, were capable of converting [Het-s*] strains into [Het-s] strains and to induce cell death with HET-S (Fig. 5, B and C; *SI Appendix*, Fig. S8). Hence, we termed this phenotype [HEC^S^]. Importantly, [HEC^Φ^] and [HEC^S^] phenotypes remained stable in time and faithfully self-propagated over several passages (*SI Appendix*, Table S7, Table S8). In addition, we observed that GFP-HEC aggregates from [HEC^S^] strains partially co-localized with [Het-s] and not with [Φ], while fluorescent HEC aggregates from [HEC^Φ^] strain produced the opposite results and co-localized predominantly with [Φ] (*SI Appendix*, Fig. S8). We concluded that the engineered HEC sequence could propagate as two distinct prion strains [HEC^Φ^] and [HEC^S^] and was capable of breaching the cross-seeding barrier between [Het-s] and [Φ] prions. Because HEC was designed by the targeted replacement of HRAM-specific residues, we concluded that such residues play a key role in the control of prion propagation, allowing or preventing cross-seeding between HRAM families.

### Cross-seeding induced structural plasticity of HEC near HRAM-specific residues

To pinpoint determinants of the cross-seeding at structural level, we investigated the engineered protein sequence *in vitro* using ssNMR. Based on the resonance assignment of recombinant ^13^C,^15^N-labelled sample, HEC adopts highly similar cross-β fold to HET-s and HELLF (*SI Appendix*, Fig. S9). We then assembled HEC fibrils *in vitro* in presence of 5% of unlabelled HET-s or HELLF, respectively termed HEC^S^ and HEC^Φ^. The resulting fibrillar states were analysed using solid-state NMR and the detected spectral fingerprints were similar between the three HEC preparations (HEC, HEC^S^ and HEC^Φ^), while exhibiting slight structural differences at some amino acid positions (*SI Appendix*, Fig. S10). Both co-aggregation experiments (HEC + 5% HET-s and HEC + 5% HELLF) produced relatively similar chemical shift perturbations in HEC, suggesting similar interaction surfaces between the peptides. To gain further insight into the structural differences between HEC co-aggregated with HET-s or HELLF, we compared both chemical shift sets and plotted the differences as a function of the primary sequence (Fig. 5D; *SI Appendix*, Fig. S10). We detected small chemical shift variations on residues constituting the amyloid core of HEC^S^ and HEC^Φ^, suggesting slight conformational changes within the assemblies (Fig. 5, D and E; *SI Appendix*, Fig. S10). Remarkably, two of the residues showing highest chemical shift difference between HEC^S^ and HEC^Φ^ were part of the HRAM-defining positions that were modified in HELLF to engineer HEC (Fig. 5A). The two residues were V250 and E251, replacing the HRAM5 hallmark residues F250 and G251 from the QxFG sub-motif, situated in the R2 repeat of HEC (Fig. 5). In addition, we found three other HRAM-defining positions in HEC (A223, N238 and N248) showing significant chemical shift displacements (> 0.5 ppm) during cross-seeding by both HELLF and HET-s (*SI Appendix*, Fig. S10). These results indicate that the engineered HEC sequence adopts similar amyloid fold to HET-s and HELLF, while exhibiting limited structural plasticity near several different HRAM-specific residues during cross-seeding. Although these conformational changes are subtle, they are in agreement with the observed HEC prion strains *in vivo* and highlight the role of HRAM-specific residue as potential hotspots defining prion infectivity and controlling cross-seeding.

## Discussion

Our study exploits the natural diversity occurring in a superfamily of fungal prion domains to document the limits of sequence-to-fold conservation in amyloids and the molecular and structural determinants of cross-seeding. First, by structurally characterizing highly divergent natural prion amyloids of the HRAM family, we uncovered that functional amyloids can evolve in a regime of fold conservation, withstanding extreme sequence diversification. The finding indicates that the sequence-to-fold evolutionary interplay for functional amyloids is similar to what have been described for globular (32) and membrane proteins (33). Second, our results shed light in an unprecedented way on the determinants of amyloid cross-seeding. We found that virtually identical amyloid backbone structures might not be sufficient for cross-seeding and that critical side-chain positions could determine the seeding specificity of an amyloid fold. Extrapolating our conclusions to amyloids causing neurodegeneration, especially considering the structural similarities between the HRAM β-solenoid fold and β-solenoid folds of pathological amyloids in humans (17, 18, 34, 35), provides novel conceptual boundaries for the molecular behaviour for this category of pathogenic agents.

## Materials and Methods

### Prion propagation and incompatibility assays

Incompatibility phenotypes were determined by confronting strains of solid corn meal agar medium and a ‘barrage’ reaction was assessed 2-3 days post-contact. Prion propagation was assayed as the ability to transmit the [φ] prion phenotype from a [φ]-donor strain to a [φ*] prion-free tester strain after confrontation on solid medium. Transformants were confronted to wild type strains either directly or after contact with a [φ]-donor strain 6, 11 and 17 days after transfection to evaluate [φ*] and [φ] pnenotypes frequencies and spontaneous [φ] prion propagation. Protein transfection experiments with amyloid fibrils of recombinant HELLF(209-277) or HET-s(218-289) were carried following the general protocol described by Benkemoun *et al*. (36) with minor modifications. In brief, an agar piece (~ 5 mm3) covered with fresh (24 h of growth) prion-free mycelium is placed in a 2 ml screw cap tube containing 500 μl of STC buffer (0.8 M sorbitol, 50 mM CaCl2, 100 mM Tris-HCl pH 7,5) in addition to 50 μl of amyloids (2-3 mg.ml-1). The mycelium is fragmented using a mechanical cell disruptor (FastPrepTM FP120). Two consecutive runs of 30 s each at speed of 6 m/s were realized and fractions of the suspension (20-25 μl) were directly spotted on corn meal medium to be assessed for prion conversion after 4-5 days of regeneration at 26°C.

### Protein purification

Cells were sonicated on ice in a lysis buffer (Tris 50 mM, 150 mM NaCl, pH 8) and centrifuged to remove *E. coli* contaminants. Proteins were expressed in inclusion bodies due to their insoluble properties and purified under denaturing conditions. The supernatant was discarded and the pellet incubated with lysis buffer supplemented with 2 % Triton X-100. The membrane pellet containing inclusion bodies was extensively washed with lysis buffer to remove Triton X-100 traces and incubated at 60 °C overnight with 8 M guanidine hydrochloride until complete solubilization. After a centrifugation step at 250 000 g, lysate was recovered and incubated for 2 h with pre-equilibrated Ni-NTA beads (Ni Sepharose 6 Fast Flow, GE Healthcare Life Sciences) in a binding buffer (50 mM Tris, 0.5 M NaCl, 20 mM imidazole, 7 M urea, pH 8). Proteins were eluted from the beads with 10 mL of elution buffer (50 mM Tris, 0.5 M NaCl, 500 mM imidazole, 7 M urea, pH 8). After the affinity chromatography step, proteins were loaded on a HiPrep 26/10 desalting column (GE Healthcare) to exchange buffer for 1 % acetic acid and remove low molecular weight compounds. All the purification steps were realized in 100 % H2O. It allowed amide proton back exchange under denaturing conditions for the mixed (1/1) [(U-^1^H/^14^N)/(U-^1^H_N_/^2^H/^15^N)] sample.

### Assembly of HELLF(209-273) fibrils in vitro

The pure protein recovered after HPLC purification in 1% acetic acid was concentrated in Amicon Ultra-15 (cut-off 3 kDa) centrifugal filter units (Merck Millipore) to reach the final protein concentration of 1 mM. For the mixed (1/1) [(U-^1^H/^14^N)/(U-^1^H_N_/^2^H/^15^N)] sample, monomers were solubilized in 1% acetic acid at 1 mM concentration at a molar ratio of 1:1. Fibrils were formed by adjusting the pH with 3 M Tris to a pH value of 7.5. The protein solution was allowed to self-assemble for 2 weeks at room temperature under slow shaking. Fibrils were then centrifuged at 20000 g and washed several times with water supplemented with 0.02 % NaN_3_ and transferred to the ssNMR rotor.

### Solid-state NMR spectroscopy of HELLF(209-277)

ssNMR spectra were recorded on a 23.5 T (1 GHz iH frequency) spectrometer (Bruker Biospin, Germany) equipped with a 0.7 mm triple resonance (^1^H, ^13^C, ^15^N) MAS probe. Sample spinning frequency was 100 kHz. Spectra were referenced according to 4,4-dimethyl-4-silapentane-1-sulphonic acid (DSS) signals.

### Backbone and side-chain resonances assignment of HELLF

We used a set of eight 3D ^1^H detected experiments: (HCA)CB(CA)NH [4 scans, 11 ms (t3) x 4 ms (t2) x 20 ms (t1)], (HCO)CA(CO)NH [16 scans, 11 ms (t3) x 8 ms (t2) x 20 ms (t1)], (H)CANH [4 scans, 11 ms (t3) x 8 ms (t2) x 20 ms (t1)], (H)CONH [8 scans, 11 ms (t3) x 8 ms (t2) x 20 ms (t1)], (H)CO(CA)NH [32 scans, 11 ms (t3) x 8 ms (t2) x 20 ms (t1)], (HCA)CBCAHA [4 scans, 8 ms (t3) x 4 ms (t2) x 20 ms (t1)], (H)N(CO)CAHA [8 scans, 14 ms (t3) x 6 ms (t2) x 20 ms (t1)] and (H)NCAHA [4 scans, 14 ms (t3) x 8 ms (t2) x 20 ms (t1)]. The combination of these experiments allowed the connectivities between each ^1^H-^15^N couple to intra-residual or sequential CA, CA, CO and HA resonances, necessary to perform the entire backbone assignment. Side-chain proton assignment was performed using a ^1^H-^13^C CP-based sequence followed by a WALTZ mixing step (H)CwaltzCH [2 scans, 6 ms (t3) x 6 ms (t2) x 20 ms (t1)]. We used as a starting point the previously assigned CA and HA chemical shifts to correlate unambiguously all the carbons and protons of the side chains step by step (27). Spectra were analysed using the CCPNmr Analysis Software (37).

### Collection of intra- and inter-molecular restraints for HELLF from solid-state NMR

211 distance restraints per monomer were collected from ^1^H detected ssNMR spectra to determine the 3D structure of the HELLF(209-273) amyloid fibril. We added to these restraints 34 dihedral angles (phi/psi) estimated from chemical shifts using the TALOS+ software. 143 long-range intra-molecular restraints were assigned on 3D (H)CHH [4 scans, 6 ms (t3) x 4 ms (t2) x 20 ms (t1)], HhNH [8 scans, 8 ms (t3) x 4 ms (t2) x 20 ms (t1)] and H(H)CH [4 scans, 6 ms (t3) x 4 ms (t2) x 20 ms (t1)] spectra using a radio frequency-driven recoupling (RFDR) ^4^H-^4^H mixing. 33 inter-molecular restraints were assigned on a 3D H(H)NH spectrum [48 scans, 9 ms (t3) x 4 ms (t2) x 20 ms (t1)] using a RFDR ^4^H-^4^H mixing, using a mixed (1/1) [(U-^1^H/^14^N)/(U-^1^H_N_/^2^H/^15^N)] sample for the unambiguous detection of intermolecular restraints. A RFDR mixing followed by a ^15^N-edited CP allowed a magnetization transfer from all the protons of the (U-^4^H/^14^N) HELLF monomers to ^15^N atoms of the (U-^4^H_N_/^2^H/^15^N), back protonated monomers adjacent in the fibril, followed by the ^1^H detection of the amide protons

### NMR structure calculation of HELLF

The structure of HELLF fibrils was determined in several cycles of structure calculations and restraint analysis with ARIA 2.3 (38). Cross-peak assignments for ^1^H-^1^H correlations were converted into distance restraints with an upper-bound of 8.5Å. Backbone dihedrals angles were predicted with TALOS+(30) from ^1^H, ^13^C and ^15^N chemical shifts. TALOS predictions for residues in secondary structure elements (β1-8 and C-terminal α-helix) were converted in dihedral angle restraints with an error range corresponding to ± 1.5 times the TALOS error with a minimum of ± 15°. HELLF fibrils structure was calculated as a pentamer with five copies of HELLF(220-272) using simulated annealing performed with CNS 1.2 (39). The ladder topology was maintained during the calculation through distance restraints ensuring that the distance between equivalent Cα atoms in neighboring monomers is constant throughout the pentamer, without fixing a particular distance value, i.e. d_m/m+1_ = d_m+1/m+2_ = d_m+2/m+3_ = d_m+3/m+4_ = d_m+4/m+5_ (40, 41). Additionally, an NCS restraint was added to minimize the r.m.s difference between atomic coordinates of the monomers(41). For every ARIA iteration 100 structures were calculated and the 10 lowest-energy structures from the last iteration were refined in a shell of water molecules (42). On the basis of the identified β-strands and the in/out distribution of side-chains from an initial ARIA calculation using NMR restraints only, intra- and inter-monomer hydrogen bond restraints between β-strands (β1-β5, β2-β6, β3-β7, β4-β8) were included in subsequent rounds of ARIA calculation. Final restraints and structure statistics are given in Supplementary Table 3.

### Solid-state NMR spectroscopy of HEC, HEC, HEC^S^ and HEC^Φ^ D

Proton-detected ssNMR spectra were recorded on a 23.5 T (1 GHz ^1^H frequency) and 14.1 spectrometers (Bruker Biospin, Germany) equipped with 0.7 mm, 1.3 mm and 3.2 mm triple resonance (^4^H, ^13^C, ^15^N) MAS probes. Spinning frequency was maintained at 100 kHz (0.7 mm) and 60 kHz (1.3 mm) for ^1^H detection and 11 kHz (3.2 mm) for ^13^C detection, respectively.

2D (H)NH experiments of HEC and HED were recorded at 100 kHz MAS (1 GHz ^1^H frequency spectrometer). Resonance assignment of HED, recorded using a 1.3 mm probe (1 GHz ^1^H frequency spectrometer), was performed using the following set of experiments: (H)CONH, (H)(CO)CA(CO)NH, (H)NCAH, (H)COCAH. 2D ^13^C-^43^C experiments of HEC^S^ and HEC^Φ^ were recorded at 11 kHz MAS (600 MHz ^1^H frequency spectrometer).

Experimental data were processed using TopSpin and analyzed using Sparky (43) or CCPNMR.

### Modeling of HRAMs families structures

3D models of the prion forming domain (PFD) of representative proteins of HRAM-2, HRAM-3 and HRAM-4 were constructed with the program MODELLER (44) using the HET-s(218-289) (PDB 2KJ3) (40) and HELLF(220-272) structures. For each HRAM family, the sequence of the R1 and R2 repeats of HET-s and HELLF were aligned on the predicted repeats of the PFD to be modeled. HRAMs models were built as trimers using the atomic coordinates of the R1 and R2 repeats of HELLF and HET-s structures as templates.

## Data Availability

All data from this work are included in the main text and the SI Appendix of the paper.

## Acknowledgments

We acknowledge financial support from the European Research Council (ERC) under the European Unions Horizon 2020 research and innovation programme (ERC-2015-CoG GA no. 648974 to G.P. and ERC-2015-StG GA no. 639020 to A.L.), IdEx Bordeaux (Chaire d’Installation to B.H., ANR-10-IDEX-03-02), the ANR (ANR-14-CE09-0020-01 to A.L., ANR-13-PDOC-0017-01 to B.H. and ANR-17-CE11-0035 to S.J.S) and the CNRS (IR-RMN FR3050). J.S. and L.B.A. were supported by individual MSCA incoming fellowships (REA grant agreements n°661799 “COMPLEX-FAST-MAS” and n°624918 “MEM-MAS”). A.D. was supported by the Nouvelle Aquitaine Regional Council.

## Author contributions

D.M., L.B.A., J.S., G.P., B.H., N. E.M and A.L. performed ssNMR experiments; D.M., N. E.M. G.P., B.H. and A.L. analyzed ssNMR data; B.B. performed structure calculations; D.M., M.B., A.N. expressed, purified, and polymerized HELLF fibrils for NMR; A.D., V.C., S.J.S. carried out microscopy, prion infectivity and in vivo experiments; B.K. performed x-ray diffraction experiment; J.S.W. performed STEM experiments; A.D., D.M., B.H., A.L. and S.J.S. wrote the paper; all authors discussed the results and commented on the manuscript.

## Supplementary Information

Supplementary Materials and Methods

Figures S1 to S10

Tables S1 to S8

### Supplementary Materials and Methods

#### Strains, plasmids and media

P. anserina strains used in this study were wild-type *het-s, het-S* and the strains *Δhet-s, Δhellf* (ΔPa_3_9900) and *Δhet-s Δhellf*. The *Δhellf* strain was obtained after disruption of the gene Pa_3_9900 and replacement of the corresponding ORF with the nourseothricin-resistance gene nat. The *Δhellf* or *Δhet-s Δhellf* strains were used as recipient strains for the expression of molecular fusions of full length HELLF or the HRAM5-containig region HELLF(209-277) and the GFP (green fluorescent protein) or RFP (red fluorescent protein). The strains were co-transformed with pGEM-T (Promega) derived vectors carrying the molecular constructs and a second ‘helper’ vector carrying a phleomycin-resistance gene ble. Phleomycin-resistant transformants were selected, grown for 30 h at 26°C and screened for the expression of the transgenes using a fluorescence microscope. The expression of the molecular fusions was under the control of the constitutive *P. anserina* gpd (glyceraldehyde-3-phosphate dehydrogenase) promoter. Positive transformants were backcrossed with a strain of the same genetic background as the recipient strain in order to obtain transgene-expressing homokaryons. The *hellf* gene was amplified with oligonucleotides 5’ gcttaattaaATGGATCCCCTCAGCGTTACTGC 3’ and 5’ atagatcttgctccTTTGCTGAAGAGGTTGCTA 3’ (capital letters correspond to *P. anserina* genomic DNA sequences) and cloned in front of the GFP encoding sequence using the PacI/BglII restriction enzymes to produce the pOP-hellf-GFP vector. The *hellf(209-277)* gene fragment was amplified with oligonucleotides 5’ gcttaattaaATGAAGACTCTGTCGGCGACCCG 3’ and 5’ atagatcttgctccTTTGCTGAAGAGGTTGCTA 3’ and cloned with PacI/BglII in front of the sequence encoding the RFP to produce the pOP-hellf(209-277)-RFP vector. In addition to the BglII site, a two amino acids linker (GA) was introduced between the sequences encoding hellf and GFP/RFP and is present on both vectors (pOP-hellf-GFP and pOP-hellf(209-277)-RFP). hellf(209-277) was also amplified with oligonucleotides 5’ tgcaaagcgcggccgcAAGACTCTGTCGGCGACCCG 3’ and 5’ gaaaatatGGATCCTCATTTGCTGAAGAGGTTGC 3’ and cloned behind the GFP using NotI/BamHI restriction enzymes to generate the plasmid pGB6-GFP-hellf(209-277). The molecular fusion GFP-HELLF(209-277) contains 3 aa linker encoded by the NotI restriction site. The fusion protein GFP-HELLF(209-277), in which HELLF(209-277) is situated at the C-terminus as it is in HELLF, resulted in rapid transition to the [φ] in all observed strains, unlike the fusion HELLF(209-277)-RFP, for which the [φ*] state was seemingly more stable. To investigate the co-localization/cross-seeding of HELLF and HET-s, Δhet-s Δhellf strain was transformed both with vectors expressing fluorescently tagged versions of full length or partial HELLF and pOP-Het-s-RFP vector (*het-s* fused to RFP under the gpd promoter).

HEC was amplified witholigonucleotides 5’ ggcgcggccgcAAAACCCTGTCAGCGACACG 3’ and 5’ ggcGGATCCTCATTTCGAGAACAGGTTGGAAAAC3’ and HED was amplified with oligonucleotides 5’ ggcgcggccgcAAAACCCTGTCTGCCACACG3’ and 5’ ggcGGATCCTCATTTGGAGAACAGGTTGCTG 3’andcloned downstream of the GFP using NotI/BamHI restriction enzymes to generate the plasmid pGB6-GFP-HEC andpGB6-GFP-HED.

The sequence encoding the first 35 aa of fnt1 (Pa_3_991O) was amplified with oligonucleotides 5’ atgcaaagcgcggccgcATGGCGAGGGCGGGGCAACATGTGG 3’ and 5’ gaaaatatGGATCCTCAATGGATGTGGAGAGATCCG 3’ and cloned in the pGB6-GFP-fnt1(1-35) downstream of the GFP, using NotI/BamHI restriction sites. The fnt1(1-35) gene fragment was also cloned in front of the RFP with PacI/BglII restriction sites to generate pOP-fnt1(1-35)-RFP. This vector was used to transform a *Δhet-s Δhellf* strain expressing GFP-HELLF(209-277) for our co-localization study.

Site-directed mutagenesis was performed using the QuikChange II mutagenesis kit (Agilent) and the pOP-hellf-GFP vector as template with oligonucleotides 5’ GACCCAAACAACGAAAGGTACTCTAGCGCG 3’ and 5’ CGCGCTAGAGTACCTTTCGTTGTTTGGGTC 3’ to generate the pOP-hellf(L52K)-GFP plasmid.

For heterologous expression in *E. coli*, we cloned hellf(209-277) in pET24 (Novagen) using the NdeI/XhoI restrictions sites. The gene fragment hellf(209-277) was amplified with oligonucleotides 5’ catATGAAGACTCTGTCGGCGACCCG 3’ and 5’ ctcgagTTTGCTGAAGAGGTTGCTA 3’ and cloned in front of a polyhistidine-tag to generate pET24-hellf(209-277)-6his plasmid.

#### Microscopy

*P. anserina* hyphae were inoculated on solid medium and cultivated for 24 to 72 h at 26°C. The medium was then cut out, placed on a glass slide and examined with a Leica DMRXA microscope equipped with a Micromax CCD (Princeton Instruments) controlled by the Metamorph 5.06 software (Roper Scientific). The microscope was fitted with a Leica PL APO 100X immersion lens. To observe HELLF-GFP plasma membrane re-localization, we inoculated HELLF-GFP expressing strain at a distance of 3-4 cm from a strain expressing HELLF(209-277)-RFP. The zone of confrontation between the two strains was observed hours after contact. Electron Microscopy of HELLF fibrils, was performed as previously described in Balguerie *et al*. 2003, EMBO.

#### Cellgrowth and protein expression

The gene encoding for HELLF(209-277) was cloned into a pet24a expression vector (Novagen) containing a C-terminal six histidines purification tag. The plasmid was transformed into BL21(DE3)pLys *E. coli* strain to express HELLF protein in minimal medium M9 enriched in ^15^NH4Cl (1 g/L) and ^13^C labeled carbon sources ([U-^13^C] D-glucose, 2 g/L). 1L of minimal labeled medium was inoculated with 2% (v/v) of fresh initial culture in LB medium. The bacteria were grown at 37°C while shaking at 220 rpm and expression was induced with 0.75 mM isopropyl β-D-1-thiogalactopyranoside (IPTG) at OD600 0.7-0.8. Cells were harvested after 3h of protein expression and frozen at −80 °C until purification.

#### Cell growth and protein expression in D_2_O

To obtain high protein production yields in 100 % D_2_O medium, we adapted step by step the bacteria to the labeled nitrogen/carbon sources and the replacement of H_2_O by D_2_O. *E. coli* BL21(DE3)pLys bacteria were grown at 37 °C in 1 mL of LB medium until the OD_600nm_ reaches 0.8. Cells were harvested and resuspended in 50 mL of M9 medium 100 % H_2_O supplemented in ^15^NH4Cl (1 g/L) and [U-^2^H,^12^C] D-glucose (2 g/L) until a OD_600nm_ of 0.8. Cells were harvested and resuspended in 150 mL of M9 medium 100 % D_2_O supplemented in ^15^NH4Cl (1 g/L) and [U-^2^H,^12^C] D-glucose (2 g/L) until a OD_600nm_ around 1.3. 850 mL of the same growth medium was added to the culture to have an OD_600nm_ around 0.2. Once the culture reaches an OD_600nm_ of 0.5, protein expression was induced at 30 °C by adding 0.75 mM IPTG. Cells were harvested after 22 h of protein expression and frozen at −80 °C until purification.

**Fig. S1.**
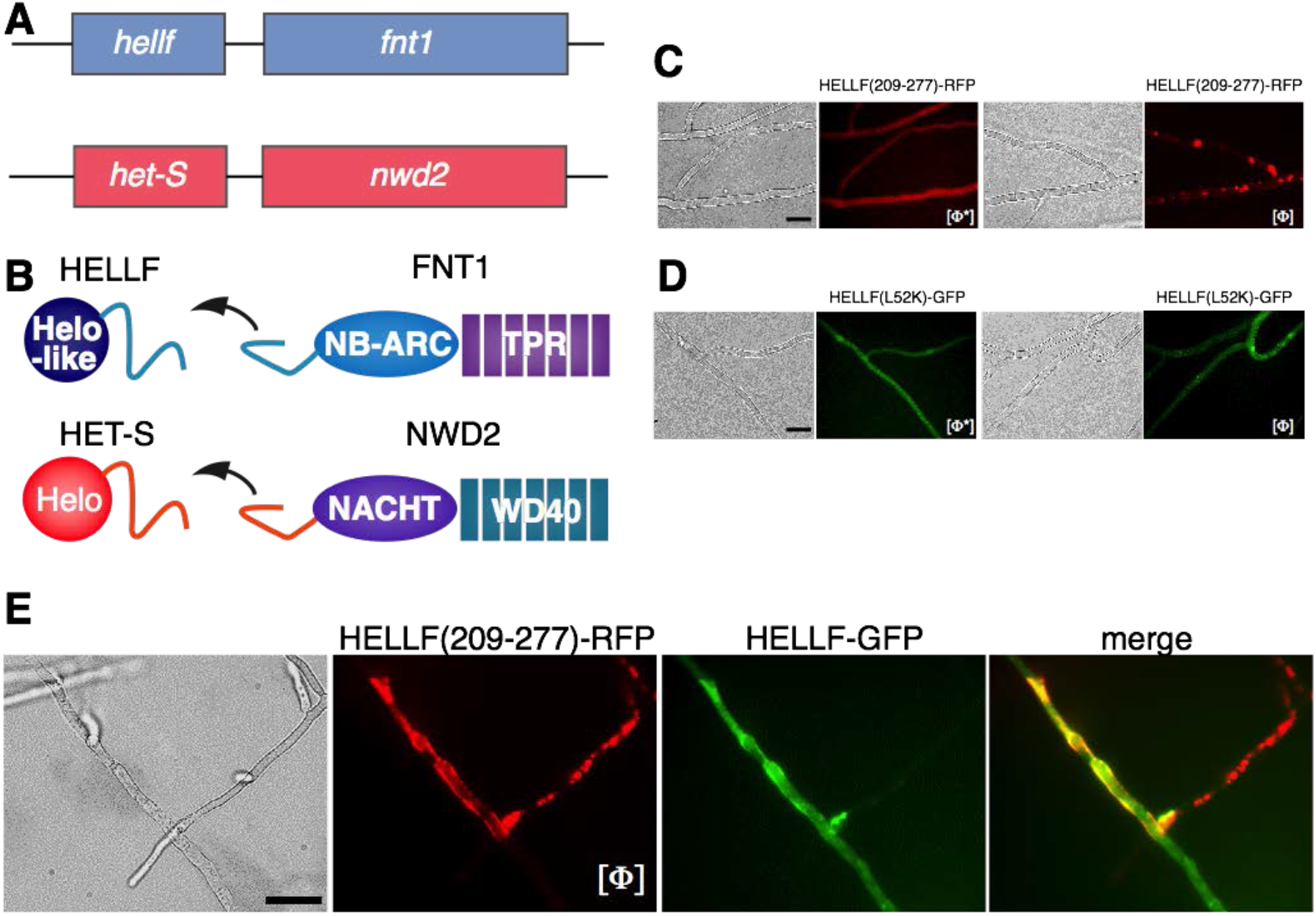
HELLF is a distant HET-S homolog. **A.** The *hellf* gene is encoded in the genome of *Podospora anserina* and is adjacent to an NLR-encoding gene termed *fnt1*. Shown is a cartoon of the gene cluster (*hellf/fnt1*), which resembles to the gene cluster formed by *het-S* and *nwd2* (*46*). **B.** Domain architecture of the cell death-inducing HELLF and HET-S proteins and the FNT1 and NWD2 NOD-like receptors. The PFD regions of HELLF and HET-S and the amyloid-forming region of the NLRs are represented by a blue and orange line, respectively. **C-D.** HELLF-based amyloid prion propagates *in vivo*. Bistability of (**C**) HELLF(209-277) and (**D**) HELLF(L52K) *in vivo*. Fluorescent microscopy images of overexpressed HELLF molecular fusions with RFP (upped panels) or GFP (lower panels), showing states of HELLF aggregation in the [Φ] strains. No aggregates are observed in [Φ*] strains. Scale bar: 5 μm. **E.** HELLF-GFP localizes in vicinity of the plasma membrane in heterokaryotic cells carrying aggregates of HELLF(209-277)-RFP (the prion [Φ] state). Scale bar: 5 μm.

**Fig. S2.**
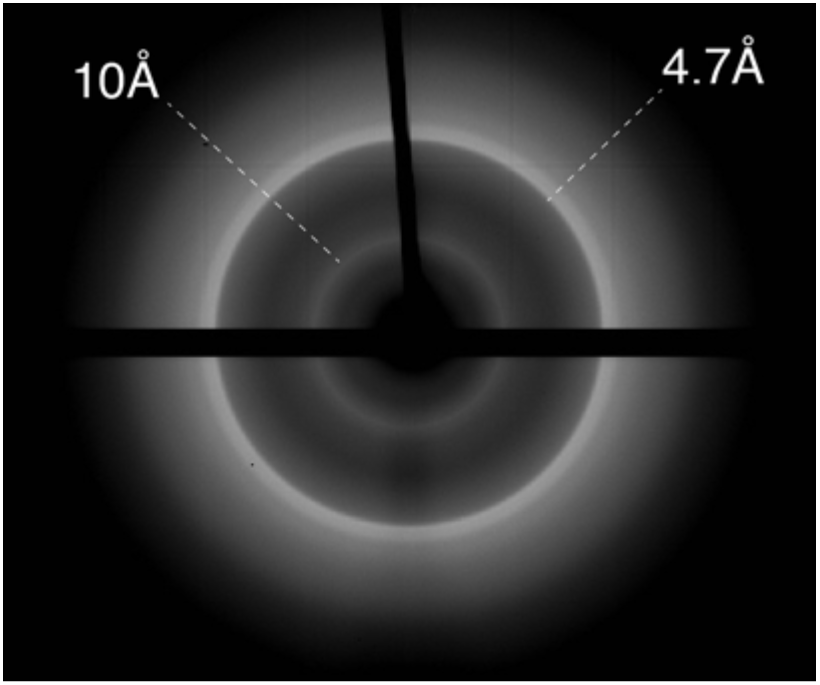
X-ray diffraction pattern of HELLF(209-277) fibrils. Major reflections appear at 4.7 and 10 Å consistent with cross-β fibrillar architecture.

**Fig. S3.**
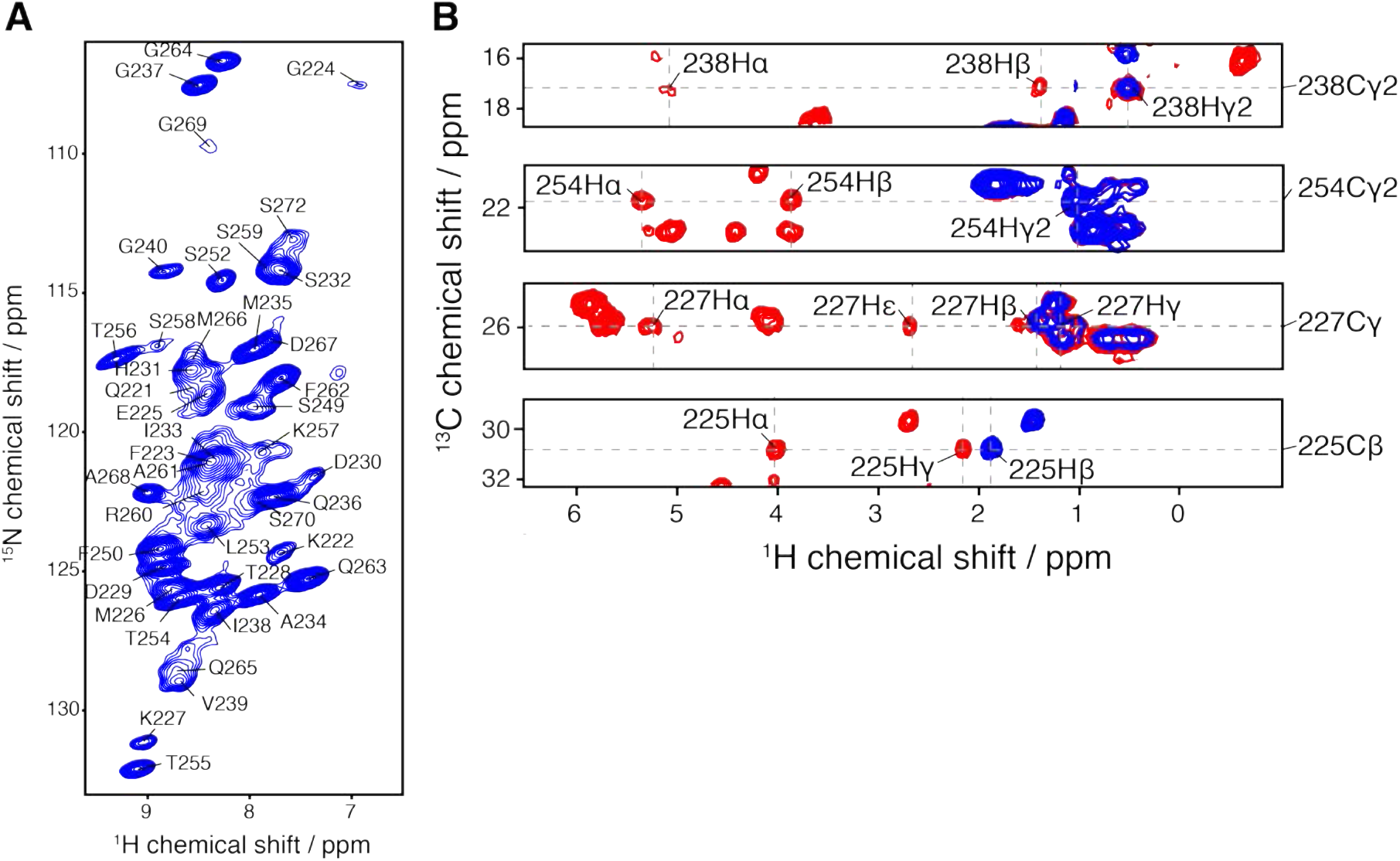
Solid-state NMR (ssNMR) characterization of HELLF(209-277) fibrils. **A.** Two-dimensional (H)NH experiment recorded on fully protonated HELLF(209-277) fibrillar assemblies. **B.** Strip plots show 13C-1H projections of the 3D (H)CCH-TOCSY experiment on HELLF(209-277) fibrillar assemblies, for selected residues.

**Fig. S4.**
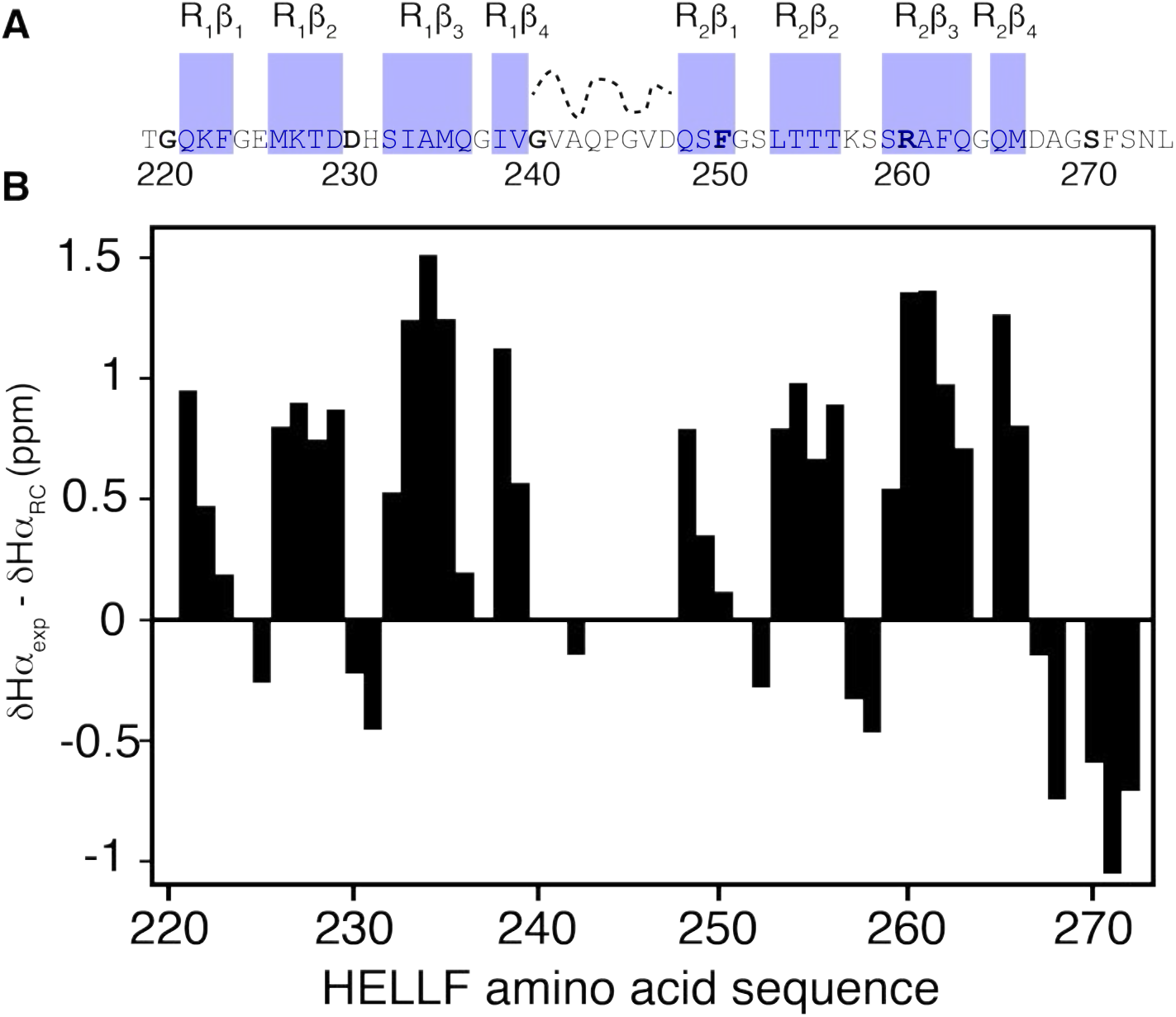
ssNMR-based secondary structure of HELLF(209-277). **A.** Beta-sheet secondary structure elements are highlighted in blue. **B.** Chemical shift index based on Hα (experimental) - Hα (random coil).

**Fig. S5.**
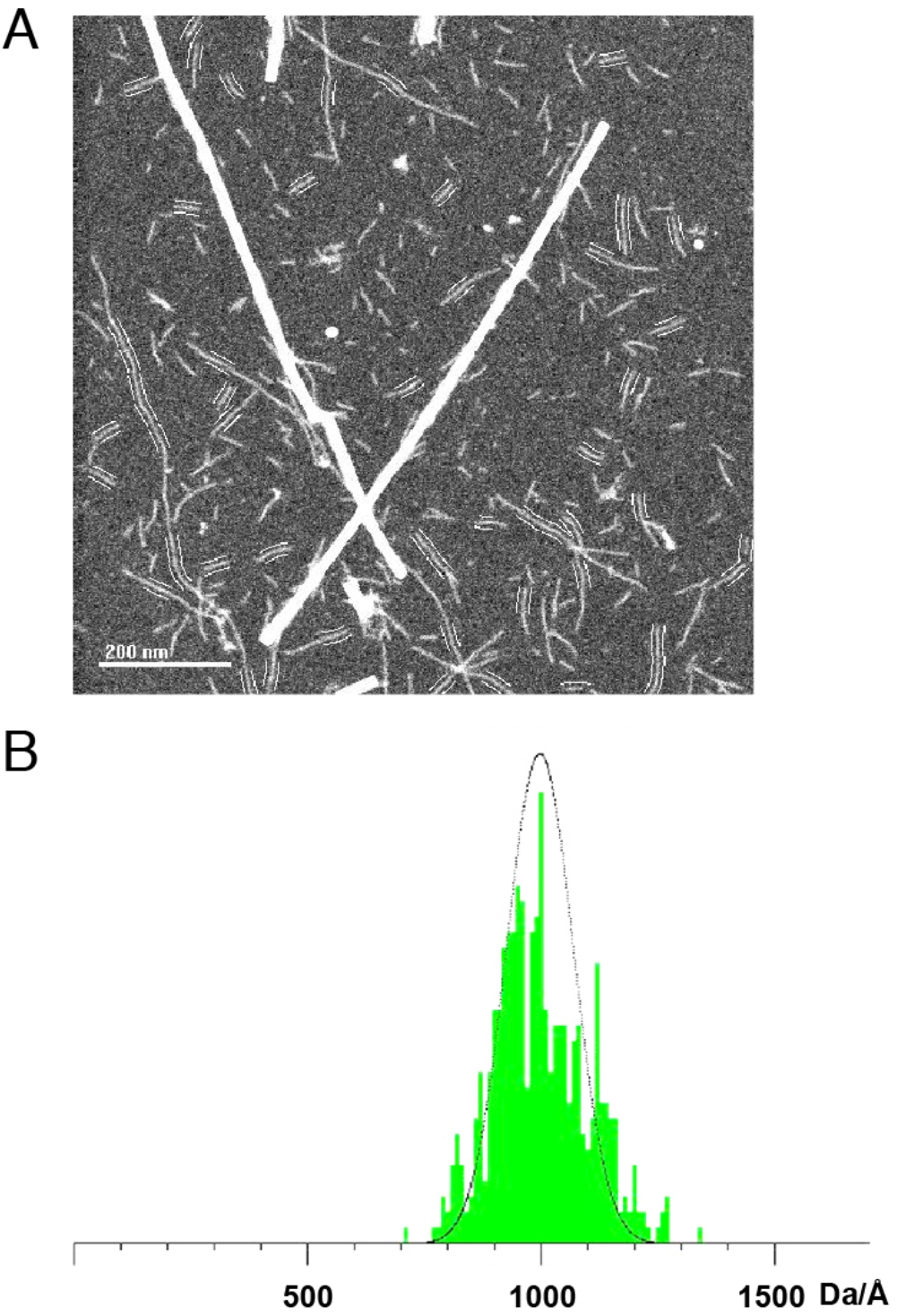
Mass-per-length measurement of HELLF(209-277) fibrillar assemblies by scanning electron transmission microscopy. **A.** Dark-field electron micrograph of HELLF (209-277) filaments and TMV reference particles. **B.** Mass-per-length histogram fitted with a Gaussian function centered at 998.533± 101.139 Da/Å.

**Fig. S6.**
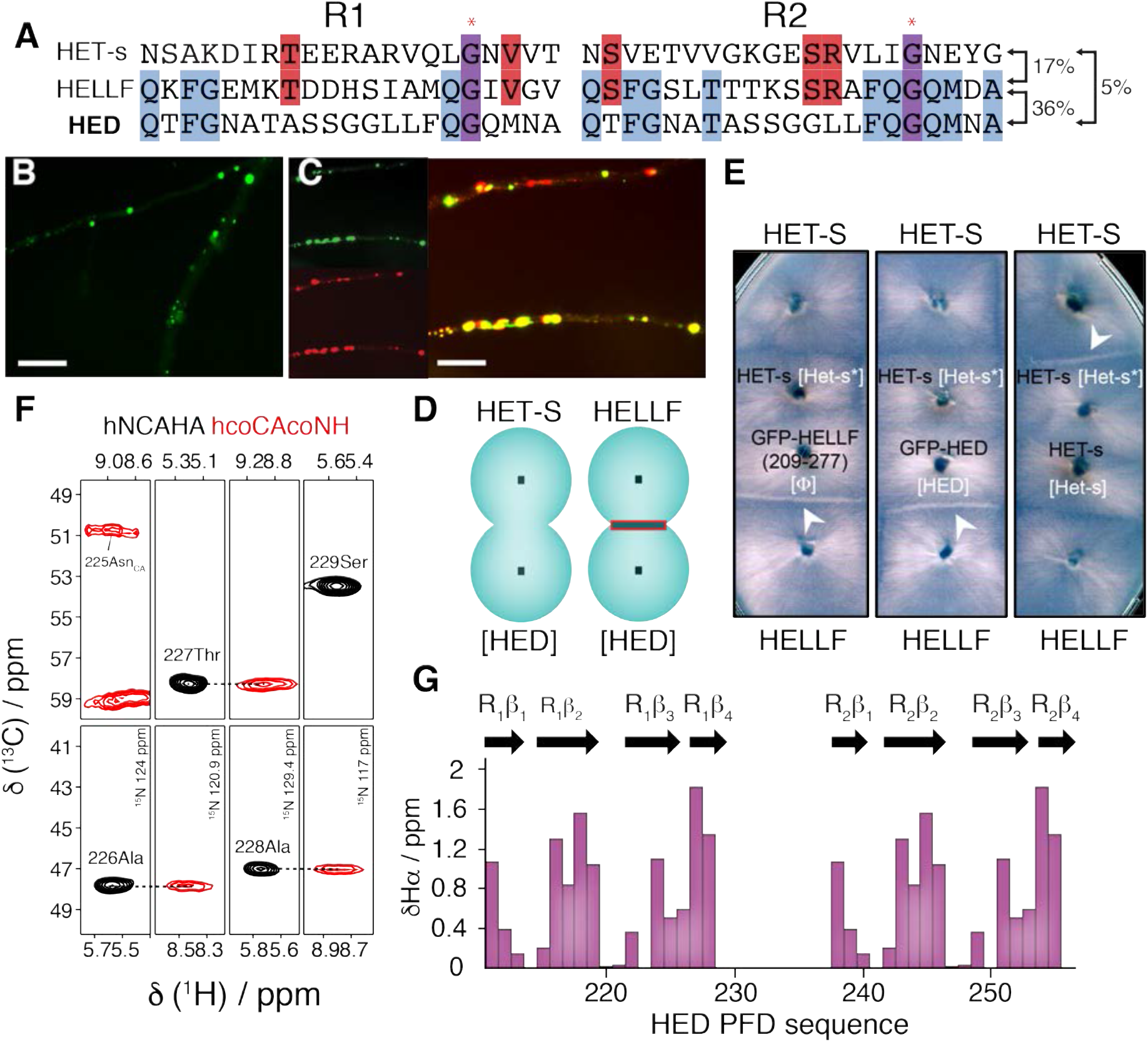
HELLF-derived protein HED sharing only 5% sequence identity with HET-s adopts a HET-s-like amyloid fold *in vitro* and can propagate as a prion *in vivo*. **A.** Sequence alignments of the two pseudo-repeats (R1 and R2), constituting the PFDs of HELLF, HET-s and the engineered protein HED. **B.** Fluorescent microscopy images of dot-like aggregates formed by HED-GFP in hyphae of *P. anserina*. Scale bar: 5 μm. **C.** Cellular co-localization (merged panel) of dot-like aggregates formed by HED-GFP (left upper panel) and HELLF(209-277)-RFP (left bottom panel). Scale bar: 5 μm. **D.** Cartoon representation of barrage phenotypes between strains expressing full-length HET-S or HELLF and strains expressing HED. The barrage reaction is shown as a line separating two incompatible strains (green circles). **E.** [HED] strains trigger programmed cell death with strains expressing full-length HELLF (lower middle panels), while being unable to induce the [Het-s] prion in a [Het-s*] strain (upper middle panels). [HED] thus behaves as [Φ] (left panels) and not like [Het-s] (right panels). Barrage reactions revealing cell death between antagonistic strains are indicated with white arrowheads. **F.** Extracts of ssNMR spectra for ^1^H, ^13^C and ^15^N sequential assignments. A combination of (HCO)CA(CO)NH (red) and (H)NCAHA (black) was used to assign HED fibrils. **G.** ssNMR-based secondary structure assignment of HED. Chemical shift index are calculated based on Hα (experimental) - Hα (random coil).

**Fig. S7.**
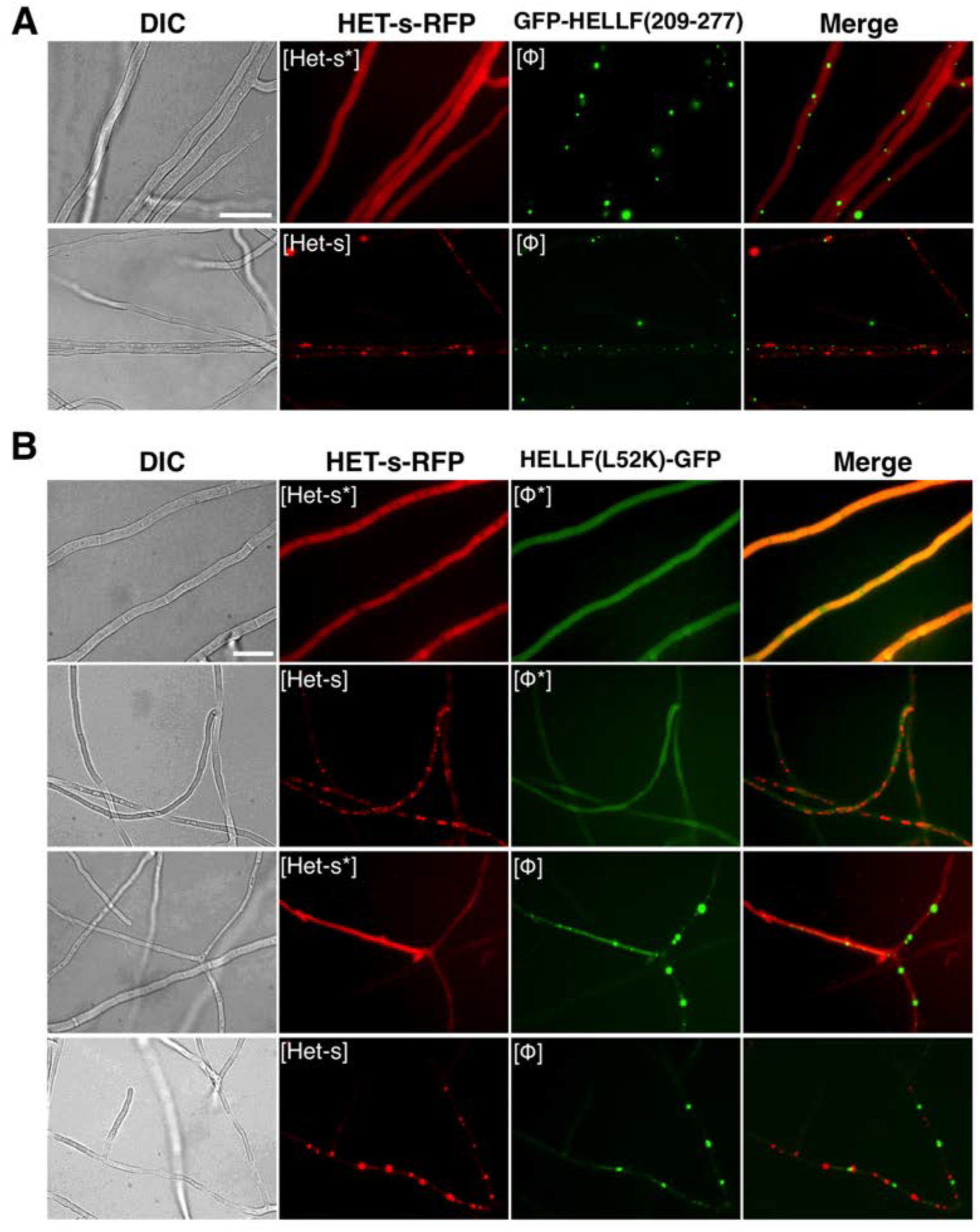
Distinct prion specificities of [Het-s] and [Φ] prions *in vivo*. **A.** Lack of cross-seeding and co-localization in vivo between [Φ] GFP-HELLF(209-277) and RFP-HET-s in soluble prion-free [Het-s*] state (top panel) or aggregated [Het-s] prion state (lower panel). Scale bar: 5 μm. **B.** Strains co-expressing HET-s-RFP ([Het-s*] or [Het-s] state) and the cytotoxic-dead HELLF(L52K)-GFP mutant ([Φ*] or [Φ] state) exhibiting all four possible epigenetic combinations ([Het-s*] [Φ*], [Het-s] [Φ*], [Het-s*] [Φ] and [Het-s] [Φ]) showing an absence of cross-seeding (and co-aggregation) of the [Φ*] state by [Het-s] prion and conversely. Co-localization was only observed when both proteins were in soluble state (top panel). Scale bar: 5 μm.

**Fig. S8.**
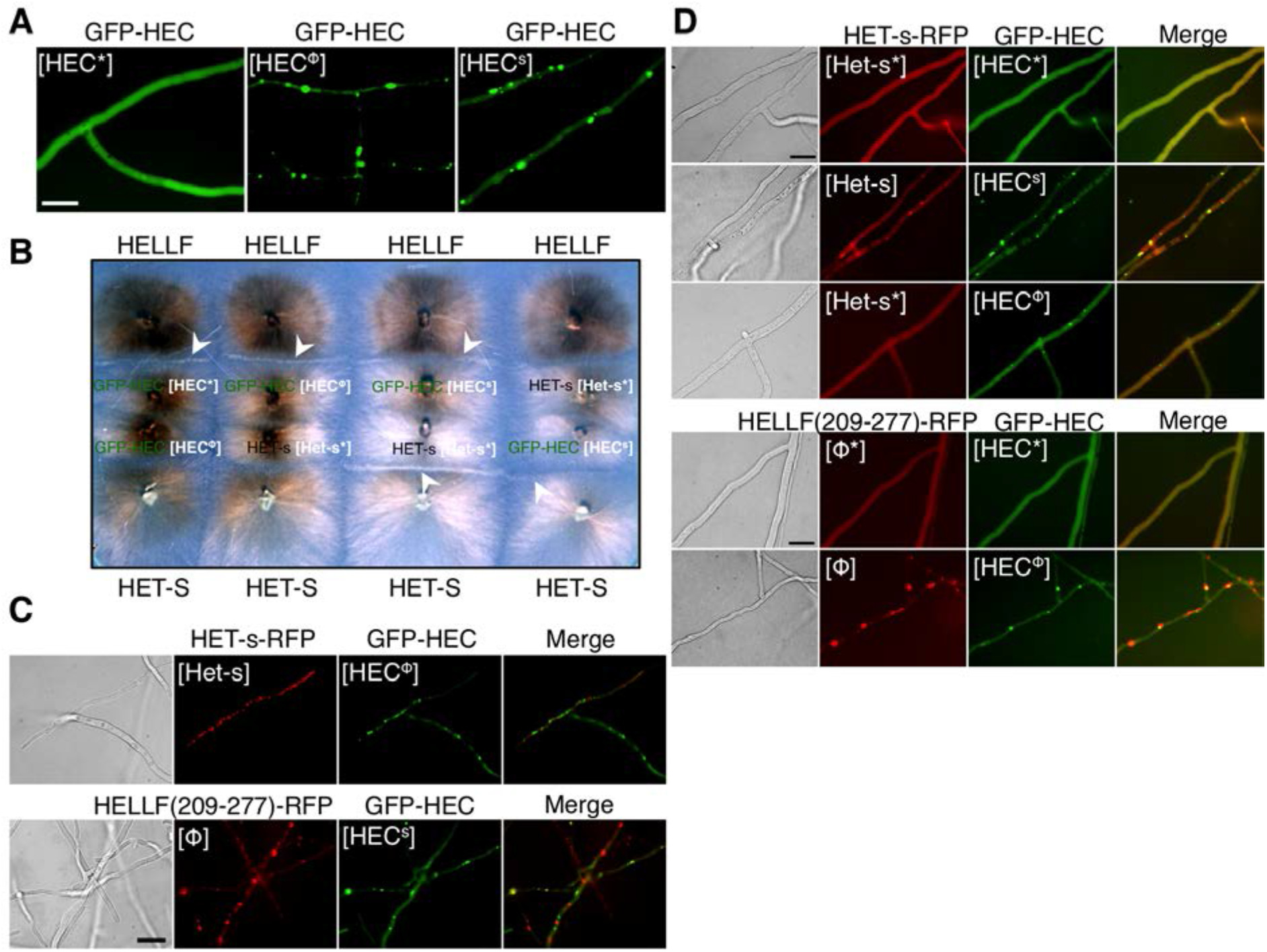
Prion strains behaviour of HEC in *P. anserina*. **A.** Fluorescent microscopy images of GFP-HEC aggregation states *in vivo*. The GFP-HEC molecular fusion shows diffuse fluorescence in [HEC*] strains (left panel) but can form dot-like fluorescent aggregates in [HEC^S^] and [HECΦ] strains after contact with prion infected [Het-s] and [φ] strains, respectively (center and right panels). Scale bar: 5μm. **B.** Barrage reactions and prion induction properties of [HEC^S^] and [HEC^φ^] strains. Two HEC-based prion strains give rise to two distinct phenotypes of the strains expressing GFP-HEC. The [HEC^φ^] strains are able to produce a barrage reaction with HELLF expressing strains but not with HET-S strain (first two tests from the left). [HEC^φ^] strains are unable to induce the [Het-s] prion in a [Het-s*] strain, while converting a [HEC*] strain into [HEC^φ^] strain. The [HEC^S^] strains produce barrage reaction with both HELLF and HET-S expressing strains (last two tests from the left). These strains are equally able to induce the [Het-s] prion in prion-free [Het-s*] strain. Barrage reactions are indicated with white arrowheads. **C.** Fluorescent microscopy images of heterokaryotic mixtures of strains expressing GFP-HEC and HET-s-RFP or HELLF(209-277)-RFP. No co-localization is observed between [Hets] and [HEC^φ^] (top row) and only a partial co-localization between [φ] and [HEC^S^] (bottom row). Scale bar: 5 μm. **D.** Fluorescent microscopy images showing co-expressions of GFP-HEC with HET-s-RFP (top three rows) or HELLF(209-277)-RFP (bottom two rows) in strains of *P. anserina*. Strains co-expressing GFP-HEC and HET-s-RFP or HELLF(209-277)-RFP present initially diffuse fluorescence for the two molecular fusions - [Het-s*][HEC*] (top row) or [φ*][HEC*] (fourth row). When these strains have been in contact with a [Het-s] strain or a [φ] strain (serving as an *in vivo* seeding strains), we observe the appearance of dot-like aggregates of GFP-HEC, which co-localize with [Het-s] for the [HEC^S^] prion strain (second row from top) and with [φ] for the [HEC^φ^] prion strain (last row from top). Prion reversion of [Het-s*][HEC*] strain by [φ] strain resulted in [Het-s*][HEC^φ^] strain (third row from top), underling the distinct prion specificity, expressed as propensity for co-localization *in vivo* with aggregates of HET-s or HELLF, of the [HEC^φ^] and [HEC^S^] HEC-based prion strains. Scale bar: 5 μm.

**Fig. S9.**
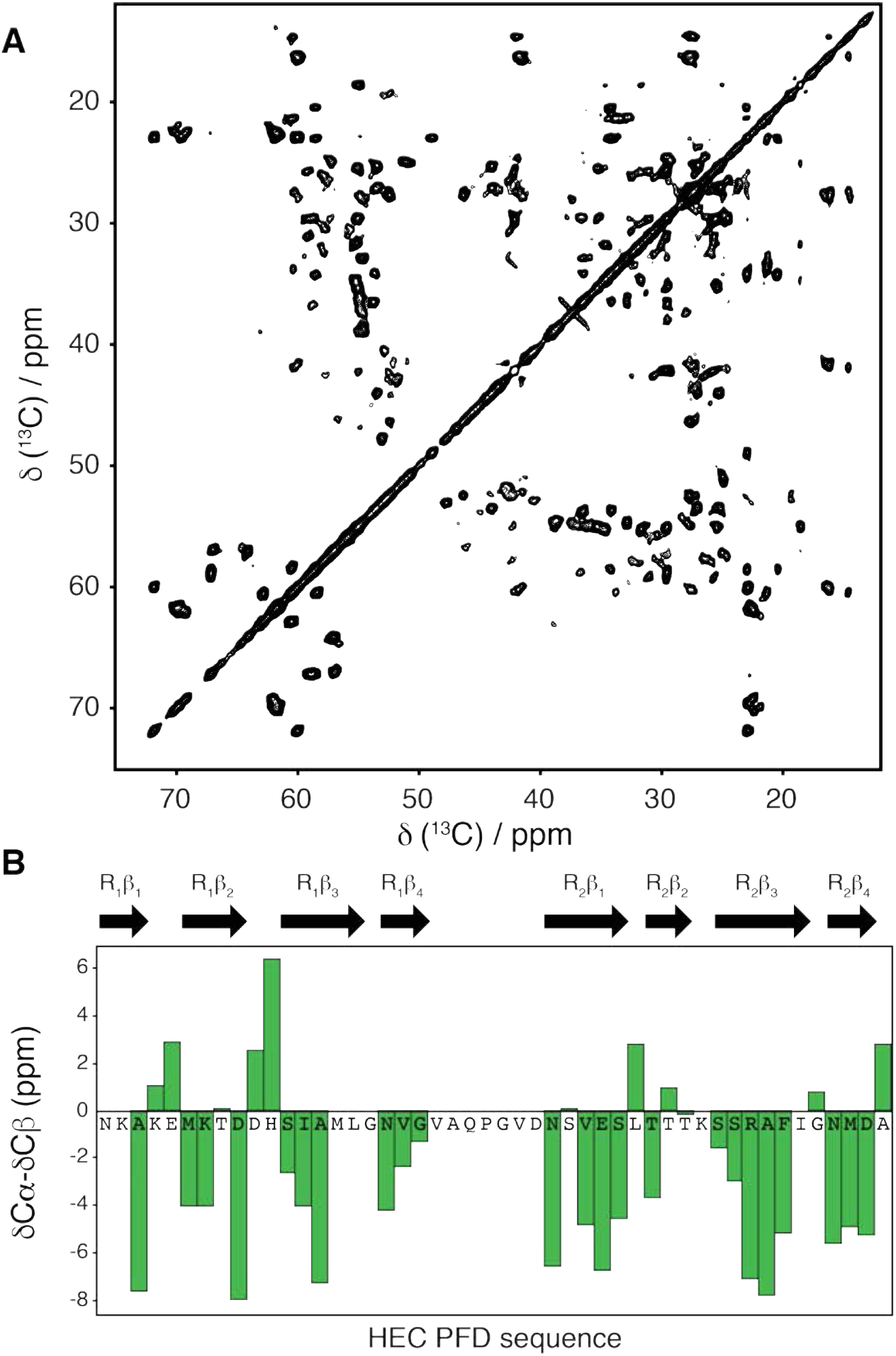
ssNMR-based secondary structure of HEC. **A.** 2D solid-state NMR ^13^C-^13^C PDSD spectrum of HEC amyloid fibrils. **B.** ssNMR-based secondary structure assignment of HEC as a function of the amino acid sequence. Negative and positive values indicate β-strand or α-helix conformation, respectively.

**Fig. S10.**
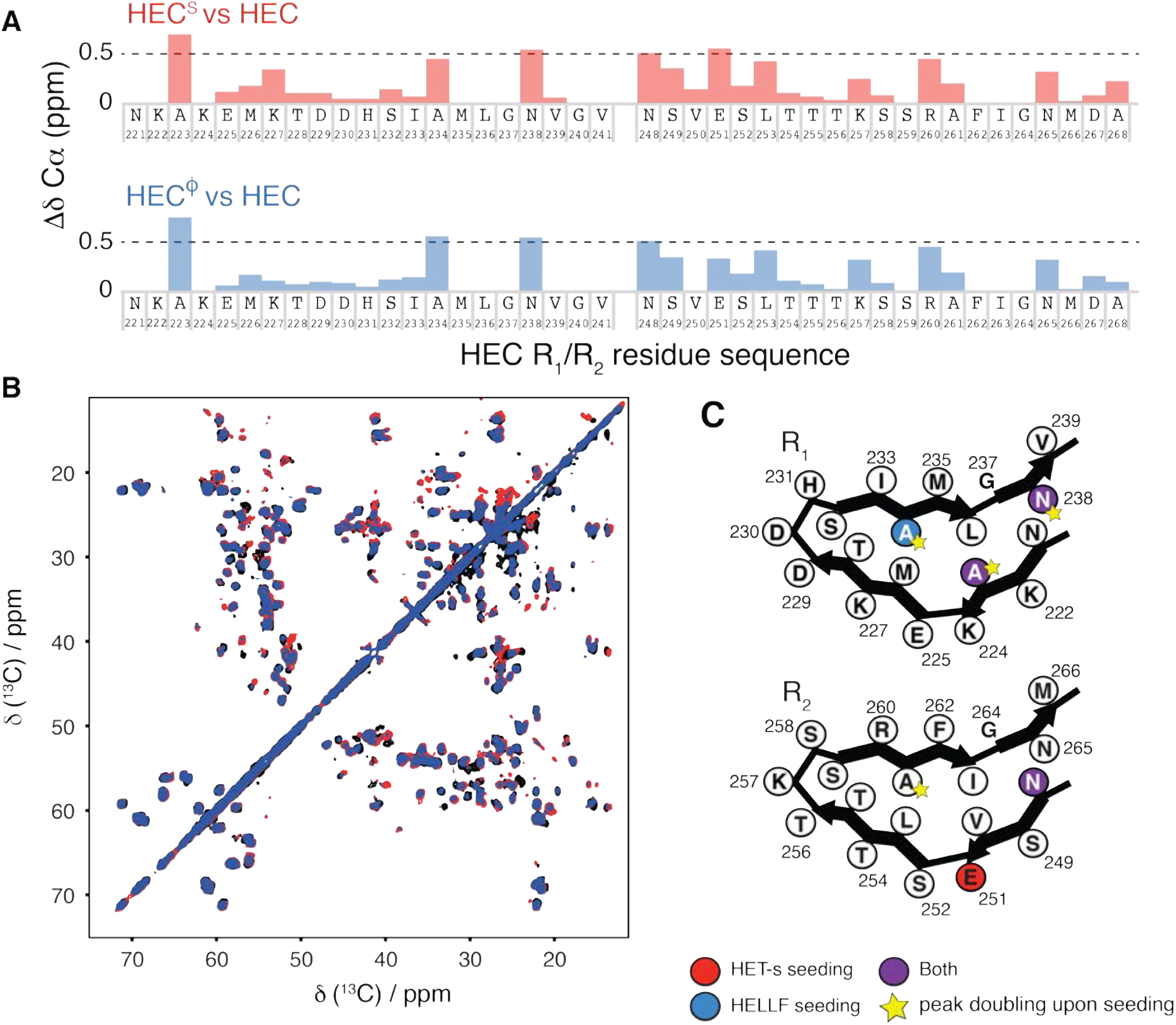
*In vitro* cross-seeding of HEC by HET-s or HELLF. **A.** Cα chemical shift differences between HEC^S^ and HEC (red) or HEC^φ^ and HEC (blue) plotted as a function of HEC primary sequence. **B.** 2D solid-state NMR ^13^C-^13^C PDSD spectra of HEC (black), HEC^S^ (red) and HECφ (blue) amyloid fibrils. **C.** Cartoon representation of the amyloid backbone of HEC pseudo-repeats R1 and R2. Residues with highest Cα chemical shift difference between HEC before and after seeding by HET-s or HELLF are highlighted in red and blue, respectively. Purple labels correspond to HEC residues affected by HET-s and HELLF cross-seeding and yellow stars highlight residues in two conformations.

**Table S1.** [Φ] prion spontaneous formation and transmission.

| recipient strain | transgene(s) | Days after transfection |  |  |  |  |  |
| --- | --- | --- | --- | --- | --- | --- | --- |
|  |  | 6 |  | 11 |  | 17 |  |
| | | [$\Phi^*$], n | [$\Phi$], n | [$\Phi^*$], n | [$\Phi$], n | [$\Phi^*$], n | [$\Phi$], n |
| *Δhellf* | HELLF(209-277)-RFP | 8 | 32 | 2 | 38 | 0 | 40 |
| *Δhellf* | HELLF(L52K)-GFP | 8 | 1 | 5 | 4 | 4 | 5 |
| *Δhellf Δhet-s* | HELLF(209-277)-RFP | 3 | 9 | 2 | 10 | 0 | 12 |
| *Δhellf Δhet-s* | HELLF(L52K)-GFP | 4 | 0 | 1 | 3 | 2 | 2 |
| *Δhellf Δhet-s* | GFP-HELLF(209-277) | 1 | 33 | 0 | 34 | nd | nd |
| *Δhellf Δhet-s* | HELLF(L52K)-GFP<br>HET-s-RFP | 8 | 0 | 5 | 3 | 4 | 4 |
| *Δhellf Δhet-s* | GFP-HELLF(209-277)<br>HET-s-RFP | 0 | 7 | nd | nd | nd | nd |

**Table S2.** Incompatibility barrage tests with strains expressing HELLF variants.

|  |  |  |
| --- | --- | --- |
| *Δhellf het-s* [Het-s] HELLF(209-277)-RFP [$\Phi^*$] | / | *hellf het-s*[Het-s] |
|  | // | *hellf het-S* |
|  | / | *Δhellf het-s* [Het-s] |
|  | // | *Δhellf het-S* |
|  | / | *Δhellf het-s* [Het-s] HELLF-GFP |
| *Δhellf het-s* [Het-s] HELLF(209-277)-RFP [$\Phi$] | / | *hellf het-s*[Het-s] |
|  | // | *hellf het-S* |
|  | / | *Δhellf het-s* [Het-s] |
|  | // | *Δhellf het-S* |
|  | / | *Δhellf het-s* [Het-s] HELLF-GFP |
| *Δhellf het-s* [Het-s] HELLF-GFP | / | *hellf het-s*[Het-s] |
|  | // | *hellf het-S* |
|  | / | *Δhellf het-s* [Het-s] |
| | // | *Δhellf het-s* [Het-s] HELLF(209-277)-RFP [$\Phi$] |
| | // | *Δhellf het-s* [Het-s] HELLF(L52K)-GFP [$\Phi$] |
| *Δhellf het-s* [Het-s] GFP-HELLF(209-277) [$\Phi$] | // | *hellf het-s*[Het-s] |
|  | // | *hellf het-S* |
|  | / | *Δhellf het-s* [Het-s] |
| *Δhellf het-s* [Het-s] HELLF(L52K)-GFP [$\Phi^*$] | / | *hellf het-s*[Het-s] |
|  | // | *hellf het-S* |
|  | / | *Δhellf het-s* [Het-s] |
| | / | *Δhellf het-s* [Het-s] HELLF(209-277)-RFP [$\Phi$] |
|  | / | *Δhellf het-s* [Het-s] HELLF-GFP |
| *Δhellf het-s* [Het-s] HELLF(L52K)-GFP [$\Phi$] | // | *hellf het-s*[Het-s] |
|  | // | *hellf het-S* |
|  | / | *Δhellf het-s* [Het-s] |
| | / | *Δhellf het-s* [Het-s] HELLF(209-277)-RFP [$\Phi$] |
|  | // | *Δhellf het-s* [Het-s] HELLF-GFP |
| *Δhellf Δhet-s* HELLF(209-277)-RFP [$\Phi^*$] | / | *hellf het-s*[Het-s] |
|  | / | *hellf het-S* |
|  | / | *Δhellf het-s* [Het-s] |
|  | / | *Δhellf Δhet-s* |
| *Δhellf Δhet-s* HELLF(209-277)-RFP [$\Phi$] | // | *hellf het-s*[Het-s] |
|  | // | *hellf het-S* |
|  | / | *Δhellf het-s* [Het-s] |
| | / | Δhellf het-S |
| | / | Δhellf Δhet-s |
| Δhellf Δhet-s GFP-HELLF(209-277) [$\Phi^*$] | / | hellf het-s[Het-s] |
|  | / | hellf het-S |
| | / | Δhellf het-s [Het-s] |
| | / | Δhellf Δhet-s |
| Δhellf Δhet-s GFP-HELLF(209-277) [$\Phi$] | // | hellf het-s[Het-s] |
|  | // | hellf het-S |
| | / | Δhellf het-s [Het-s] |
| | / | Δhellf het-S |
| | / | Δhellf Δhet-s |
| Δhellf Δhet-s HELLF(L52K)-GFP [$\Phi^*$] | / | hellf het-s[Het-s] |
|  | / | hellf het-S |
| | / | Δhellf het-s [Het-s] |
| | / | Δhellf het-S |
| | / | Δhellf Δhet-s |
| | / | Δhellf het-s [Het-s] HELLF(209-277)-RFP [$\Phi$] |
| | / | Δhellf het-s [Het-s] HELLF-GFP |
| Δhellf Δhet-s HELLF(L52K)-GFP [$\Phi$] | // | hellf het-s[Het-s] |
|  | // | hellf het-S |
| | / | Δhellf het-s [Het-s] |
| | / | Δhellf het-S |
| | / | Δhellf Δhet-s |
| | / | Δhellf het-s [Het-s] HELLF(209-277)-RFP [$\Phi$] |
| | // | Δhellf het-s [Het-s] HELLF-GFP |
| Δhellf Δhet-s HELLF(L52K)-GFP [$\Phi^*$]<br>HET-s-RFP [Het-s*] | / | hellf het-s[Het-s] |
|  | / | hellf het-S |
| | / | Δhellf het-s [Het-s] |
| | / | Δhellf het-S |
| | / | Δhellf Δhet-s |
| | / | Δhellf het-s [Het-s] HELLF(209-277)-RFP [$\Phi$] |
| | / | Δhellf het-s [Het-s] HELLF-GFP |
| Δhellf Δhet-s HELLF(L52K)-GFP [$\Phi^*$]<br>HET-s-RFP [Het-s] | / | hellf het-s[Het-s] |
|  | // | hellf het-S |
| | / | Δhellf het-s [Het-s] |
| | // | Δhellf het-S |
| | / | Δhellf Δhet-s |
| | / | Δhellf het-s [Het-s] HELLF(209-277)-RFP [$\Phi$] |
| | / | Δhellf het-s [Het-s] HELLF-GFP |
| Δhellf Δhet-s HELLF(L52K)-GFP [$\Phi$]<br>HET-s-RFP [Het-s*] | // | hellf het-s[Het-s] |
|  | // | hellf het-S |
| | / | Δhellf het-s [Het-s] |
| | / | Δhellf het-S |
| | / | Δhellf Δhet-s |
| | / | Δhellf het-s [Het-s] HELLF(209-277)-RFP [$\Phi$] |
| | // | Δhellf het-s [Het-s] HELLF-GFP |
| Δhellf Δhet-s HELLF(L52K)-GFP [$\Phi$]<br>HET-s-RFP [Het-s] | // | hellf het-s[Het-s] |
|  | // | hellf het-S |
| | / | Δhellf het-s [Het-s] |
| | // | Δhellf het-S |
| | / | Δhellf Δhet-s |
| | / | Δhellf het-s [Het-s] HELLF(209-277)-RFP [$\Phi$] |
| | // | Δhellf het-s [Het-s] HELLF-GFP |
| Δhellf Δhet-s GFP-HELLF(209-277) [$\Phi$]<br>HET-s-RFP [Het-s*] | // | hellf het-s[Het-s] |
|  | // | hellf het-S |
| | / | Δhellf het-s [Het-s] |
| | / | Δhellf het-S |
| | / | Δhellf Δhet-s |
| | // | Δhellf het-s [Het-s] HELLF-GFP |
| Δhellf Δhet-s GFP-HELLF(209-277) [$\Phi$]<br>HET-s-RFP [Het-s] | // | hellf het-s[Het-s] |
|  | // | hellf het-S |
| | / | Δhellf het-s [Het-s] |
| | // | Δhellf het-S |
| | / | Δhellf Δhet-s |
| | // | Δhellf het-s [Het-s] |

A barrage incompatibility reaction is marked with as //, a normal contact as /

**Table S3.**
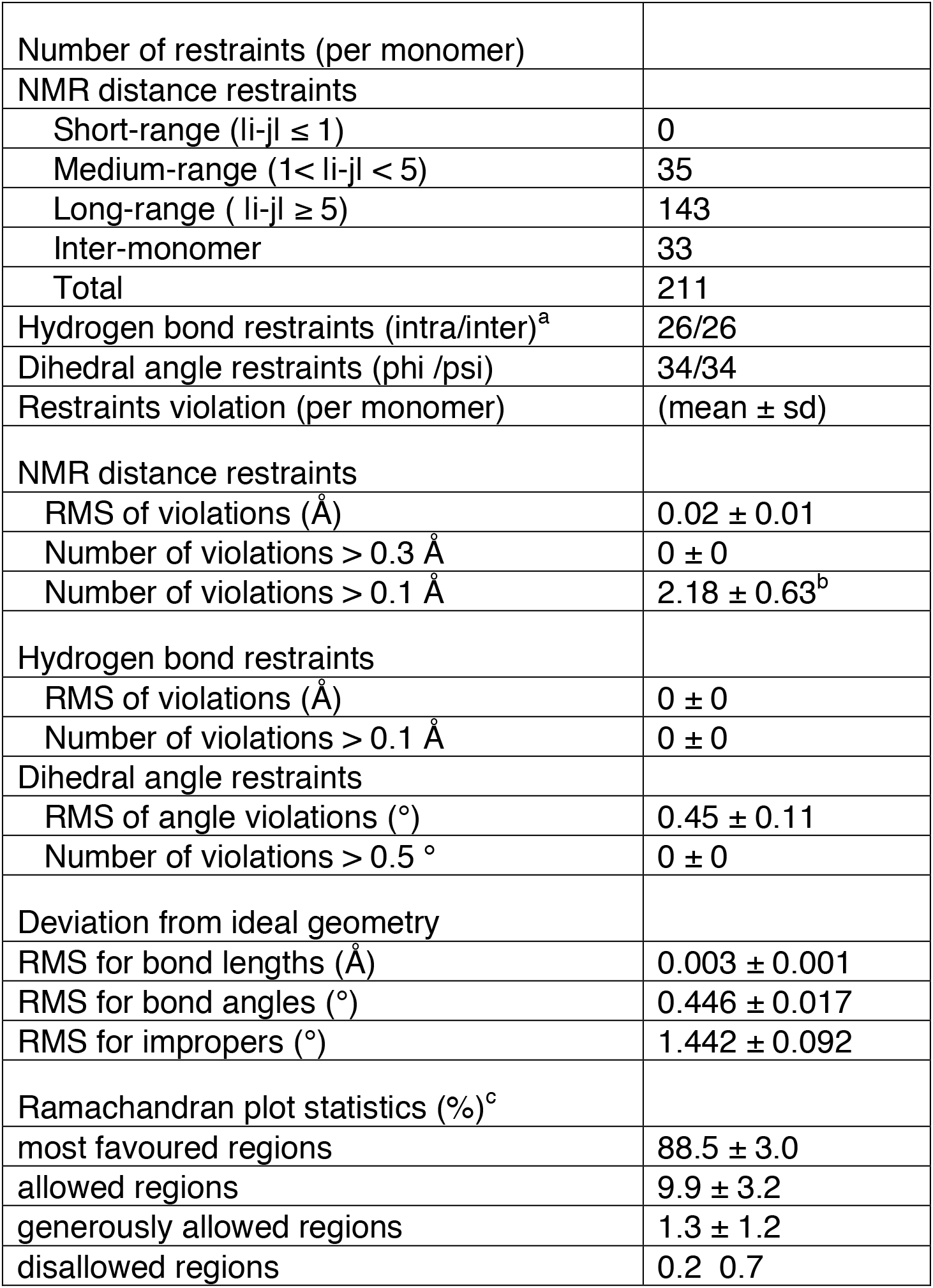
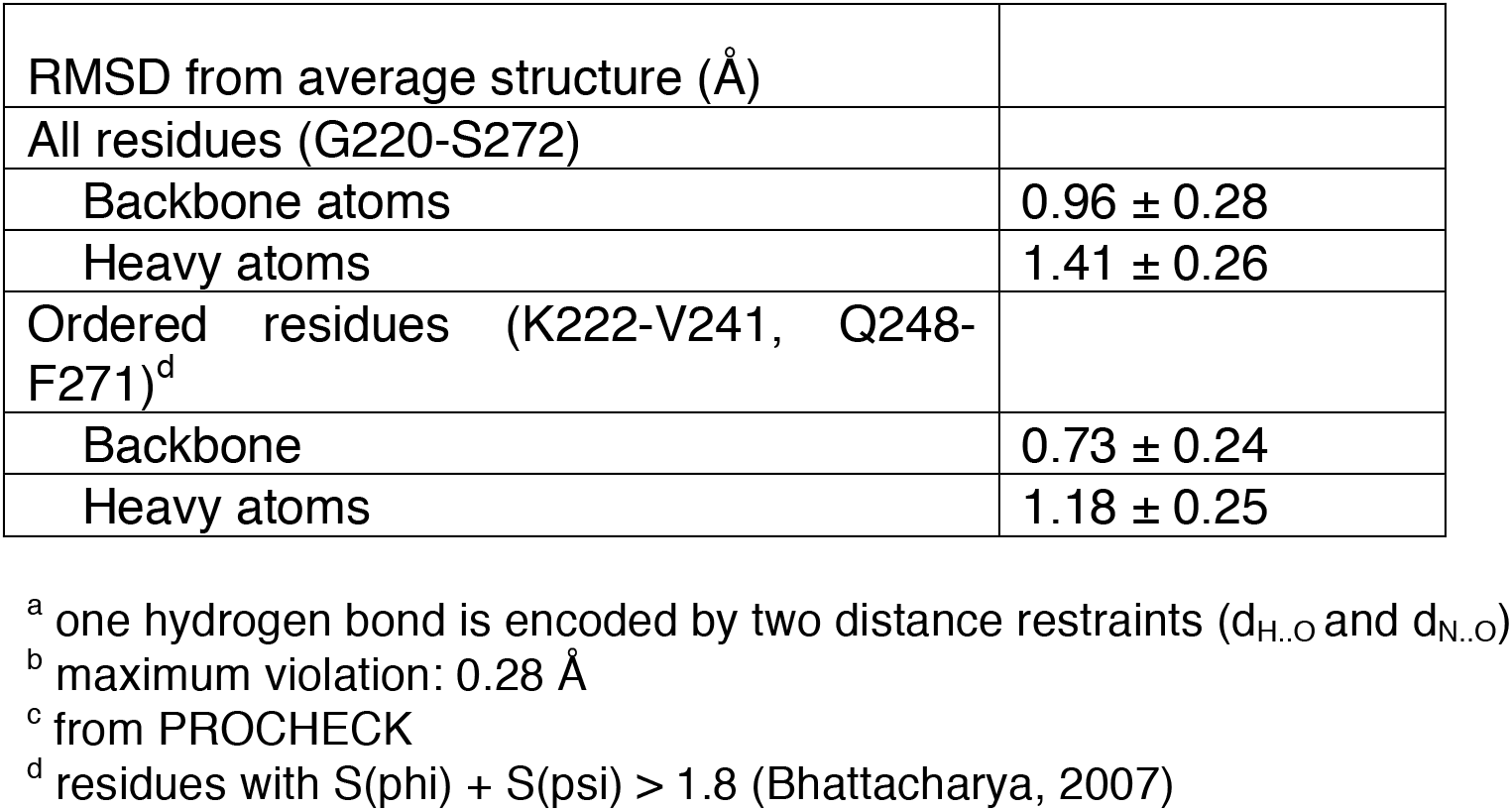
NMR structure calculation statistics.

**Table S4.** Transfection of HET-s and HELLF fibrils.

| recipient strain | no protein | HET-s(218-289) fibrils | HELLF(209-277) fibrils |
| --- | --- | --- | --- |
| *het-s* ([Het-s]/[Het-s*]) | 0/28 | 26/28 | 0/28 |
| *hellf*(209-277) ([Φ]/[Φ*]) | 1/40 | 0/20 | 6/52 |

**Table S5.**
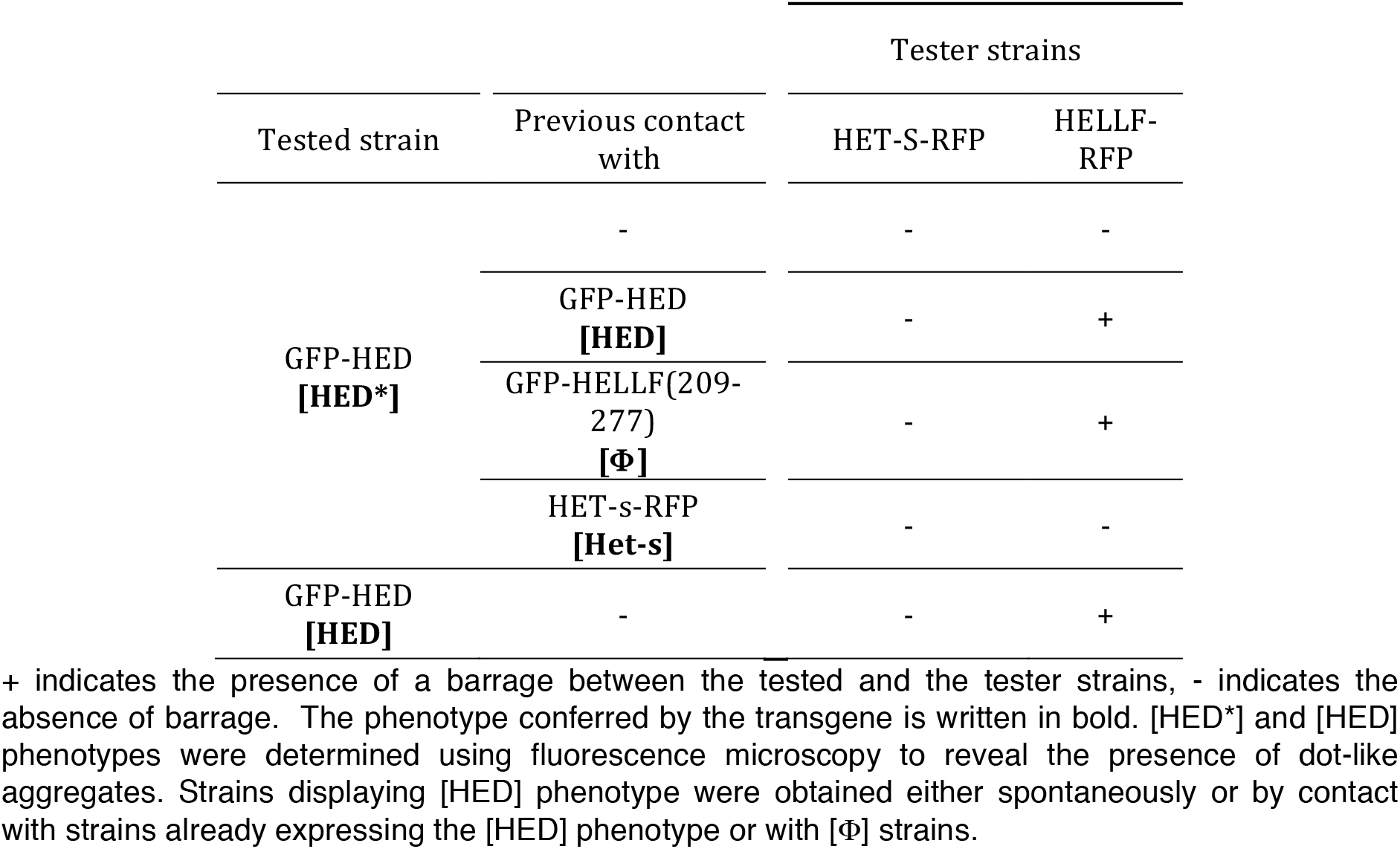
[HED] prion transmission and activation of HELLF cell death activity.

**Table S6.** [HED] and [HEC^Φ^] prion spontaneous formation and transmission.

| Transgene | days after transfection |  |  |  |
| --- | --- | --- | --- | --- |
|  | 4 |  | 8 |  |
|  | [HED*]<br>(n) | [HED]<br>(n) | [HED*]<br>(n) | [HED]<br>(n) |
| GFP-HED | 6 | 19 | 0 | 19 |
|  | 4 |  | 230 |  |
|  | [HEC*]<br>(n) | [HEC•]<br>(n) | [HEC*]<br>(n) | [HEC•]<br>(n) |
| GFP-HEC | 26 | 5 | 142 | 16 |

**Table S7.** [HEC^S^] and [HEC^Φ^] prions transmission and incompatibility phenotype.

| Tested strain | Previous contact with | Tester strains |  |
| --- | --- | --- | --- |
|  |  | HET-S-RFP | HELLF-RFP |
| GFP-HEC<br>[HEC*] | - | - | - |
|  | GFP-HELLF(209-277)<br>[Φ] | - | + |
|  | HET-s-RFP<br>[Het-s] | +* | + |
|  | GFP-HEC<br>[HEC<sup>S</sup>] | +* | + |
|  | GFP-HEC<br>[HEC•] | - | + |
| GFP-HEC<br>[HEC<sup>S</sup>] | - | +* | + |
| GFP-HEC<br>[HEC•] | - | - | + |

+ indicates the presence of a barrage between the tested and the tester strains, +\* indicates the presence of an attenuated barrage, - indicates the absence of barrage. The phenotype conferred by the transgene is written in bold. [HEC\*], [HEC•] and [HEC<sup>S</sup>] phenotypes were determined using fluorescence microscopy to reveal the presence of dot-like aggregates and on the ability to induce [Het-s\*] to [Het-s] conversion. Strains displaying [HEC<sup>S</sup>] phenotype were obtained by contact with [Het-s] strains or with strains already expressing the [HEC<sup>S</sup>] phenotype. Strains displaying [HEC•] phenotype were obtained either spontaneously or by contact with [Φ] strains or with strains already expressing the [HEC•] phenotype.

**Table S8.**
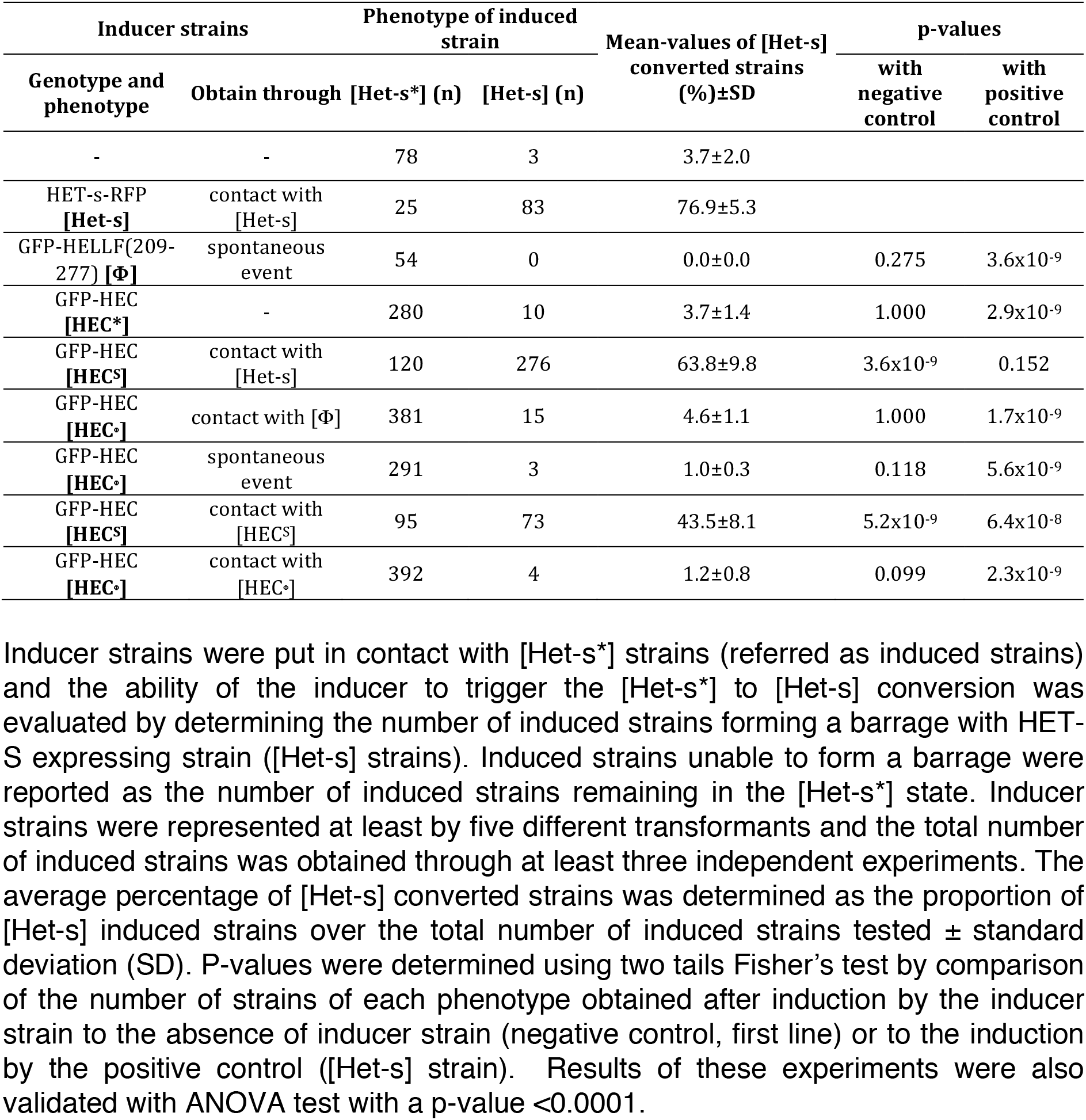
Monitoring of the ability of GFP-HEC transformants to convert [Het-s^*^] strain to [Het-s] phenotype.

## Bibliography

1. Chiti F, Dobson CM (2017) Protein misfolding, amyloid formation, and human disease: A summary of progress over the last decade. Annu Rev Biochem 86:27–68.

2. Prusiner SB (2013) Biology and genetics of prions causing neurodegeneration. Annu Rev Genet 47:601–623.

3. Eisenberg D, Jucker M (2012) The amyloid state of proteins in human diseases. Cell 148(6):1188–1203.

4. Watts JC, Prusiner SB (2018) β-Amyloid Prions and the Pathobiology of Alzheimer’s Disease. Cold Spring Harb Perspect Med 8(5).

5. Prusiner SB (2012) Cell biology. A unifying role for prions in neurodegenerative diseases. Science 336(6088):1511–1513.

6. Sanders DW, et al. (2014) Distinct tau prion strains propagate in cells and mice and define different tauopathies. Neuron 82(6):1271–1288.

7. Rey NL, et al. (2019) α-Synuclein conformational strains spread, seed and target neuronal cells differentially after injection into the olfactory bulb. Acta Neuropathol Commun 7(1):221.

8. Collinge J, Clarke AR (2007) A general model of prion strains and their pathogenicity. Science 318(5852):930–936.

9. Qiang W, Yau W-M, Lu J-X, Collinge J, Tycko R (2017) Structural variation in amyloid-β fibrils from Alzheimer’s disease clinical subtypes. Nature 541(7636):217–221.

10. Morales R, Moreno-Gonzalez I, Soto C (2013) Cross-seeding of misfolded proteins: implications for etiology and pathogenesis of protein misfolding diseases. PLoS Pathog 9(9):e1003537.

11. Ren B, et al. (2019) Fundamentals of cross-seeding of amyloid proteins: an introduction. J Mater Chem B, Mater Biol Med 7(46):7267–7282.

12. Sampson TR, et al. (2020) A gut bacterial amyloid promotes α-synuclein aggregation and motor impairment in mice. Elife 9.

13. Horvath I, Wittung-Stafshede P (2016) Cross-talk between amyloidogenic proteins in type-2 diabetes and Parkinson’s disease. Proc Natl Acad Sci USA 113(44):12473–12477.

14. Lim KH (2019) Diverse Misfolded Conformational Strains and Cross-seeding of Misfolded Proteins Implicated in Neurodegenerative Diseases. Front Mol Neurosci 12:158.

15. Ritter C, et al. (2005) Correlation of structural elements and infectivity of the HET-s prion. Nature 435(7043):844–848.

16. Wasmer C, et al. (2008) Amyloid fibrils of the HET-s(218-289) prion form a beta solenoid with a triangular hydrophobic core. Science 319(5869):1523–1526.

17. Vázquez-Fernández E, et al. (2016) The structural architecture of an infectious mammalian prion using electron cryomicroscopy. PLoS Pathog 12(9):e1005835.

18. Fitzpatrick AWP, et al. (2017) Cryo-EM structures of tau filaments from Alzheimer’s disease. Nature 547(7662):185–19O.

19. Daskalov A, et al. (2015) Signal transduction by a fungal NOD-like receptor based on propagation of a prion amyloid fold. PLoS Biol 13(2):e1002O59.

20. Riek R, Saupe SJ (2016) The HET-S/s Prion Motif in the Control of Programmed Cell Death. Cold Spring Harb Perspect Biol 8(9).

21. Saupe SJ (2011) The [Het-s] prion of Podospora anserina and its role in heterokaryon incompatibility. Semin Cell Dev Biol 22(5):46O–468.

22. Seuring C, et al. (2012) The mechanism of toxicity in HET-S/HET-s prion incompatibility. PLoS Biol 10(12):e1001451.

23. Greenwald J, et al. (2010) The mechanism of prion inhibition by HET-S. Mol Cell 38(6):889–899.

24. Loquet A, Saupe SJ (2017) Diversity of amyloid motifs in NLR signaling in fungi. Biomolecules 7(2).

25. Daskalov A, Dyrka W, Saupe SJ (2015) Theme and variations: evolutionary diversification of the HET-s functional amyloid motif. Sci Rep 5:12494.

26. Agarwal V, et al. (2014) De novo 3D structure determination from sub-milligram protein samples by solid-state 100 kHz MAS NMR spectroscopy. Angew Chem Int Ed Engl 53(45):12253–12256.

27. Andreas LB, et al. (2016) Structure of fully protonated proteins by proton-detected magic-angle spinning NMR. Proc Natl Acad Sci USA 113(33):9187–9192.

28. Samoson A (2019) H-MAS. J Magn Reson 306:167–172.

29. Stanek J, et al. (2016) NMR Spectroscopic Assignment of Backbone and Side-Chain Protons in Fully Protonated Proteins: Microcrystals, Sedimented Assemblies, and Amyloid Fibrils. Angew Chem Int Ed Engl 55(50):15504–15509.

30. Shen Y, Delaglio F, Cornilescu G, Bax A (2009) TALOS+: a hybrid method for predicting protein backbone torsion angles from NMR chemical shifts. J Biomol NMR 44(4):213–223.

31. Loquet A, Giller K, Becker S, Lange A (2010) Supramolecular interactions probed by 13C-13C solid-state NMR spectroscopy. J Am Chem Soc 132(43):15164–15166.

32. Rost B (1999) Twilight zone of protein sequence alignments. Protein Engineering Design and Selection 12(2):85–94.

33. Olivella M, Gonzalez A, Pardo L, Deupi X (2013) Relation between sequence and structure in membrane proteins. Bioinformatics 29(13):1589–1592.

34. Falcon B, et al. (2018) Structures of filaments from Pick’s disease reveal a novel tau protein fold. Nature 561(7721):137–140.

35. Falcon B, et al. (2019) Novel tau filament fold in chronic traumatic encephalopathy encloses hydrophobic molecules. Nature 568(7752):420–423.

36. Benkemoun L, et al. (2006) Methods for the in vivo and in vitro analysis of [Het-s] prion infectivity. Methods 39(1):61–67.

37. Stevens TJ, et al. (2011) A software framework for analysing solid-state MAS NMR data. J Biomol NMR 51(4):437–447.

38. Bardiaux B, Malliavin T, Nilges M (2012) ARIA for solution and solid-state NMR. Methods Mol Biol 831:453–483.

39. Brunger AT (2007) Version 1.2 of the Crystallography and NMR system. Nat Protoc 2(11):2728–2733.

40. Van Melckebeke H, et al. (2010) Atomic-resolution three-dimensional structure of HET-s(218-289) amyloid fibrils by solid-state NMR spectroscopy. J Am Chem Soc 132(39):13765–13775.

41. Nilges M (1993) A calculation strategy for the structure determination of symmetric dimers by 1H NMR. Proteins 17(3):297–309.

42. Linge JP, Williams MA, Spronk CAEM, Bonvin AMJJ, Nilges M (2003) Refinement of protein structures in explicit solvent. Proteins 50(3):496–506.

43. Lee W, Tonelli M, Markley JL (2015) NMRFAM-SPARKY: enhanced software for biomolecular NMR spectroscopy. Bioinformatics 31(8):1325–1327.

44. Sali A, Blundell TL (1993) Comparative protein modelling by satisfaction of spatial restraints. J Mol Biol 234(3):779–815.

